# Evaluation of noninvasive biospecimens for transcriptome studies

**DOI:** 10.1101/2022.09.06.506813

**Authors:** Molly Martorella, Silva Kasela, Renee Garcia-Flores, Alper Gokden, Stephane E. Castel, Tuuli Lappalainen

**Author notes:** Corresponding authors: Molly Martorella, Stephane Castel, Tuuli Lappalainen. These authors contributed equally to this work (Stephane E. Castel, Tuuli Lappalainen).

## Abstract

Transcriptome studies disentangle functional mechanisms of gene expression regulation and may elucidate the underlying biology of disease processes. However, the types of tissues currently collected typically assay a single post-mortem timepoint or are limited to investigating cell types found in blood. Noninvasive tissues may improve disease-relevant discovery by enabling more complex longitudinal study designs, by capturing different and potentially more applicable cell types, and by increasing sample sizes due to reduced collection costs and possible higher enrollment from vulnerable populations. Here, we develop methods for sampling noninvasive biospecimens, investigate their performance across commercial and in-house library preparations, characterize their biology, and assess the feasibility of using noninvasive tissues in a multitude of transcriptomic applications. We collected buccal swabs, hair follicles, saliva, and urine cell pellets from 19 individuals over three to four timepoints, for a total of 300 unique biological samples, which we then prepared with replicates across three library preparations, for a final tally of 472 transcriptomes. Of the four tissues we studied, we found hair follicles and urine cell pellets to be most promising due to the consistency of sample quality, the cell types and expression profiles we observed, and their performance in disease-relevant applications. This is the first study to thoroughly delineate biological and technical features of noninvasive samples and demonstrate their use in a wide array of transcriptomic and clinical analyses. We anticipate future use of these biospecimens will facilitate discovery and development of clinical applications.

## INTRODUCTION

Large scale transcriptomics has multiple applications in biomedical research. It provides insight into the functional consequences of genetic variation and aids our understanding of genetic determinants of disease mechanisms. Prospective applications include biomarker identification of disease risk, onset, prognosis, and treatment response, discovery of potential therapies, and assessing outcomes of *in vitro* perturbations of environmental or pharmacological exposures^1^. In the realm of cancer research, this approach has been fruitful in identifying early diagnostic markers, classifying cancer subtypes, identifying novel drug targets, and optimizing treatment choice^2–4^. In cases of rare, genetic disease, diagnosis is enhanced by inclusion of transcriptomic data due to improved detection and annotation of rare functional variants^5–7^. For common traits and diseases, discovery remains complicated by trait polygenicity, linkage disequilibrium, small effect sizes of variants, and widespread pleiotropy^8–11^. By integrating transcriptomic data with genetic information, loci with an effect on gene expression, known as molecular quantitative trait loci (molQTLs), lend potential interpretation to variants found by genome wide association studies (GWAS)^12–14^.

Despite their promise, there are several barriers to transcriptome studies. Recent works suggest that elucidation of key pathways and molecular targets for a given trait requires transcriptomes from relevant tissues, i.e. cell types^15, 16^, and contexts^17^. Dynamic changes in the transcriptome and molQTLs, which change over the course of development, disease, or environmental conditions, are suspected to lend even greater insight into the genetic architecture of the genome and are essential for using the transcriptome as a biomarker. At this time, whole blood and peripheral blood mononuclear cells (PBMCs) are the most readily collected tissue type and can be longitudinally sampled. However, the Genotype-Tissue Expression (GTEx) project, the largest and most comprehensive study of genetic regulation across post-mortem human tissues, showed that blood is an outlier in its gene expression regulatory mechanisms relative to other tissues of the body^18, 19^, and the majority of expression quantitative trait loci (eQTLs) in linkage disequilibrium with GWAS variants are not found in whole blood^20^. Previous observations suggest this difference is driven by the unique functions and cell type composition of blood versus other tissues of the body^15, 16, 21, 22^. This biological difference poses major limitations to discovery in contexts where the cell types in blood are not centrally implicated in trait development or disease pathogenesis^18–20^. Sampling directly from meaningful tissue types, if the tissues of interest are known, provides more biologically relevant data. However, current approaches include surgical extractions, invasive biopsies, and post-mortem donations, which cannot be sampled over time and typically result in high cost and complicated logistics^23^. These factors also contribute to underrepresentation of minorities and vulnerable populations in most transcriptomic studies^24–27^.

Thus, future transcriptome studies would benefit from sampling techniques that capture a broad array of cell types relevant to disease, enable use of longitudinal study designs for observation of context-specific effects, and facilitate rapid expansion to underserved populations to mitigate impending health disparities. Here, we investigate the use of noninvasive biospecimens to overcome these barriers. Early studies suggest buccal swabs, hair follicles, nasal swabs, saliva, and urine cell pellets may have potential use in clinical settings and for discovery^28–38^, though no studies to date have comprehensively delineated sources of technical and biological variance in noninvasive tissue types, nor their performance across a wide array of transcriptomic and clinical analyses. Furthermore, no comparisons to invasive sample types have been made in terms of cell type composition or regulation of gene expression and splicing. In this study, we collected buccal swabs, hair follicles, saliva, and urine cell pellets from 19 individuals over 4 timepoints, and we prepared the samples for sequencing using both in-house and commercial library kits. Both the sample collection and low-cost library preparation methods used here are available open-access. We investigated the unique biology of noninvasive sample types and analyzed sources of technical variance, identified suitable invasive tissue type proxies, and demonstrated their use in transcriptomic and disease-relevant applications. We conclude that hair follicles and urine cell pellets are promising biospecimens for future study.

## RESULTS

### Noninvasive tissues are amenable to low-input library preparations

Buccal swabs, hair follicles, saliva, and urine cell pellets were collected from 19 healthy individuals at three to four separate time points, amounting to 300 unique biological samples (Figure 1a). Briefly, participants deposited a saliva sample into an Oragene saliva collection kit, 10 hair follicles were plucked, and participants provided a buccal swab sample (see Methods and DOI: dx.doi.org/10.17504/protocols.io.kqdg3pjzzl25/v1). Saliva samples were stored according to kit instructions, and hair follicles and buccal swabs were flash frozen at -80C. Saliva, hair follicles, and buccal swabs were all able to be collected and stored within 9 minutes, on average. Urine samples were obtained at any time throughout the day, and underwent serial centrifugation before flash freezing the cell pellet at -80C. Individuals enrolled in the study provided genotyping data from 23andMe or Ancestry SNP arrays, or from low-pass whole genome sequencing provided by Gencove. Using the 1000 genomes database and k-nearest neighbors we find 15/19 individuals are of European descent, 3/19 are admixed Americans, and 1/19 is of East Asian ancestry.

**Figure 1.**
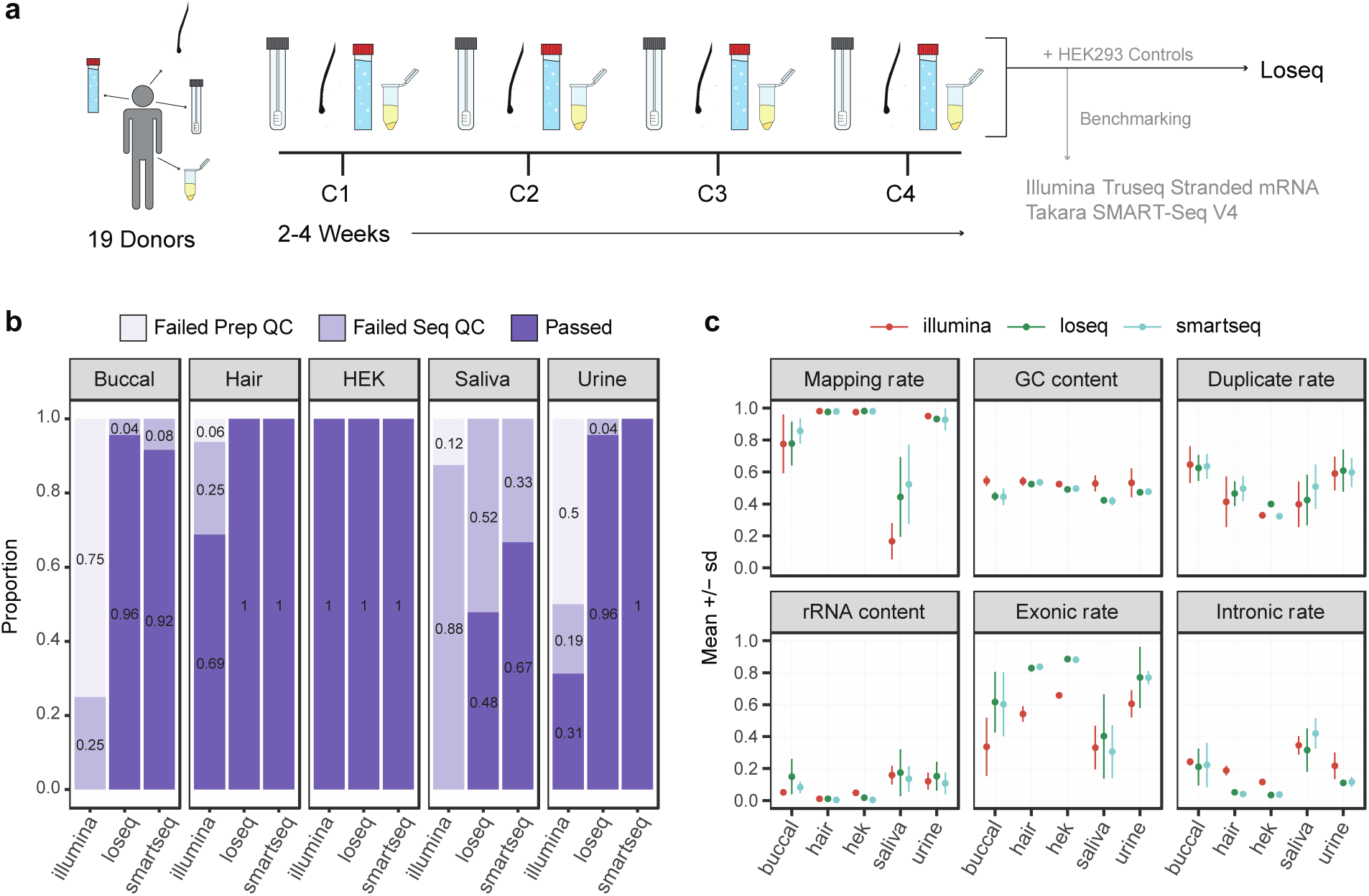
Noninvasive sample study design and processing outcomes. **a.** Four collections (C1-C4) of four noninvasive tissues were collected from 19 donors over the course of 2-4 weeks per donor. All samples were processed using our in-house method, Loseq, while a subset was prepared using commercially available kits. Two biological replicates of HEK293 cell controls were included in triplicate for all library preparations. **b.** Proportion of samples passing per tissue type and preparation. Failed Prep QC = exceeded 600bp average size or less than 2nM yield. Failed Seq QC = protein-coding and lncRNA depth less than 1 million. **c.** RNA-seq quality metrics for all sequenced samples for each tissue and library preparation.

Total RNA yielded from noninvasive tissues was tissue-specific in yield and variability (buccal median = 1.05ug and IQR = 0.56ug, hair median = 1.44ug and IQR = 1.92ug, saliva median = 6.82ug and IQR = 8.60, urine median = 0.25ug and IQR = 1.54ug, Supp. Fig. 1c). Urine pellets (Levene P = 0.0002), hair follicles (P = 0.01) and buccal swabs (P=0.05) exhibited significant donor-dependent variability in yield compared to other tissues (Supp. Fig. 1e). This observation agrees with clinically known interindividual differences in cell numbers found in urine^29^, while hair follicle output appeared to be due to individual differences in hair texture (i.e. follicles more consistently remained attached to thicker, coarser hair). All samples were prepared for sequencing using a low-cost in-house library preparation we developed specifically for low-input bulk RNA applications, which uses template-switching oligo (TSO) and tagmentation chemistry and reduces cost by 83% and 68% per reaction compared to the Illumina TruSeq Stranded mRNA Library Prep and SMART-seq V4 kits, respectively (Supp. Table 1, Supp. Table 2). We herein refer to this method as Loseq.

To validate the consistency of our in-house method, we included 15 technical replicates per tissue in each Loseq library preparation batch. To compare performance across library preparation methods, 12 randomly selected samples of each tissue were prepared using the TakaraBio Smartseq V4 kit, a commercially available kit with similar chemistry to Loseq, and 16 samples were prepared using the Illumina TruSeq Stranded mRNA kit, one of the most frequently used kits in the transcriptomics field (Supp. Fig. 1a). For all preparation methods, 2 HEK293 cell samples were included in triplicate to serve as a quality control standard. In total, 472 noninvasive and 36 HEK293 transcriptomes were prepared across the three library preps. 485 of 508 samples passed pre-sequencing quality criteria of > 2nM concentration and average library size < 600 bp and were sequenced on the Illumina NovaSeq S4 platform to a mean depth of 25.6 million total reads per sample (Supp. Fig. 1f, 1g).

Post-sequencing, we evaluated multiple quality control metrics returned from RNA-SeQC^39^. Ultimately, we found protein-coding and lncRNA read depth corresponded with high quality samples and consistent gene expression capture (Supp. Fig. 1b, Supp. Fig. 2). Across all tissues, Illumina prepared samples yielded a lower number of reads, and we thus used a less stringent 1 million protein-coding and lncRNA depth threshold for cross-preparation comparisons. A 2.5 million threshold and a 5 million threshold were used for the Loseq-only and GTEx comparison analyses, respectively.

Looking across tissues and library preparations, we observe 400 of the 508 samples pass pre- and post-sequencing quality checks (Figure 1b). Notably, hair performs well regardless of kit used and meets pre- and post-sequencing standards typical of traditionally used bulk RNA-sequencing samples (hair mean RIN = 8.9, Figure 1c, Supp. Fig. 1, Supp. Fig. 2). 8 of 16 urine and 12 of 16 buccal samples fail pre-sequencing checks for the Illumina kit. Given urine shows excellent performance via traditional RNA-sequencing quality metrics when low-input library preparation protocols are used (86/90 and 12/12 of samples pass QC for Loseq and Smartseq, respectively), we suspect this is driven by low and highly variable RNA yield not amenable to traditional bulk kits (median = 253.4ng, Q1 = 98ng, Q3 = 1642.2ng, Supp. Fig. 1c, Supp. Fig. 1e). Buccal and saliva display higher rates of RNA degradation, as reflected by lower RIN (buccal mean RIN = 2.5, saliva mean RIN = 2.6) and computationally derived transcript integrity (buccal mean TIN = 39, saliva mean TIN = 30), as well as diminished performance apparent in other RNA-seq QC metrics (Fig. 1c, Supp. Fig. 1b, Supp. Fig. 2). The lower quality of these tissue types is likely resulting in the higher rate of failure across the different preparations. We found saliva to be a particularly poor biospecimen for transcriptomic study with few samples passing our thresholds (0/16 Illumina, 43/90 Loseq, and 8/12 of Smartseq pass QC). Looking across saliva collections we observe some donors more consistently pass or fail quality checkpoints (10 donors fail 3 or more collections, 8 donors fail 1 or fewer collections, Supp. Fig. 1a), suggesting donor-specific variables may play a larger role in determining the sample quality relative to other tissue types. Buccal, though not an ideal sample type (as shown by the metrics in Figure 1c), performs well enough for use in targeted applications (86/90 and 11/12 pass QC for Loseq and Smartseq, respectively).

To compare gene expression patterns across libraries in an unbiased manner, we downsampled all QC-passed samples to a depth of 1 million by performing binomial sampling 5 times on the raw, unfiltered counts matrix and then taking the average. We found the majority of genes expressed in a tissue were captured regardless of preparation method (Supp. Fig. 3a) and there was high agreement in gene expression levels across library preparations (Supp. Fig. 3d). One difference of note is Loseq tends to capture longer genes with lower GC content relative to the commercial kits (Supp. Fig. 3b, Supp. Fig. 3c). Principal component analysis (PCA) shows that the preparation method contributed minimally to variance observed across the samples (Supp. Fig. 4).

### Unmapped reads capture tissue-specific microbial signatures

Since we observed a low mapping rate for buccal and saliva and because the noninvasive samples were collected from non-sterile human tissues, we decided to investigate biological and technical sources of unmapped reads in our samples (Supp. Fig. 5a). First, we remapped unmapped reads to microbial genomes using Decontaminer^40^. This process remapped only a small fraction of the unmapped reads (Figure 2a, Supp. Fig. 5b), a somewhat unsurprising result given our library preparation is targeted to capture poly-A mRNA transcripts and most microbial transcripts are not polyadenylated. Nonetheless, when we look at the top 0.5% most abundant remapped species we observe distinct microbial signatures across the noninvasive tissues that support previously known microbiota of the oral cavity, human skin, and genitourinary tract (Figure 2b)^41–44^. In addition, we see high correlation in estimated species abundances across technical replicates suggesting that, despite the limitations in our library preparation approach, we are capturing real, replicable biological signal (Supp. Fig. 5c). Altogether, these findings support noninvasive samples may bear biological utility in follow-up microbiome studies using microbiome-specific or total RNA library preparations. Next, we used FastQC to identify overrepresented sequences in the remaining unmapped reads. These sequences were compiled across all tissues and samples into a list of 707 sequences, which were compared to primer and adapter sequences. From this analysis we observe 517 of the highly abundant reads were present in all tissues across all library preparations (“Repeated” in Fig. 2a, Supp. Fig. 5d). This suggests these reads are most likely technical byproducts from the sequencing process, but they may also be genomic regions that align poorly. Buccal and saliva display a greater proportion of reads of unknown origin, which may result from biological sources, like microbial species that our analysis did not identify, or from technical factors, like sample degradation leading to short sequences, overamplification, and poor alignment. Overall, especially for saliva and for many buccal samples, a large proportion of reads remained of unknown origin.

**Figure 2.**
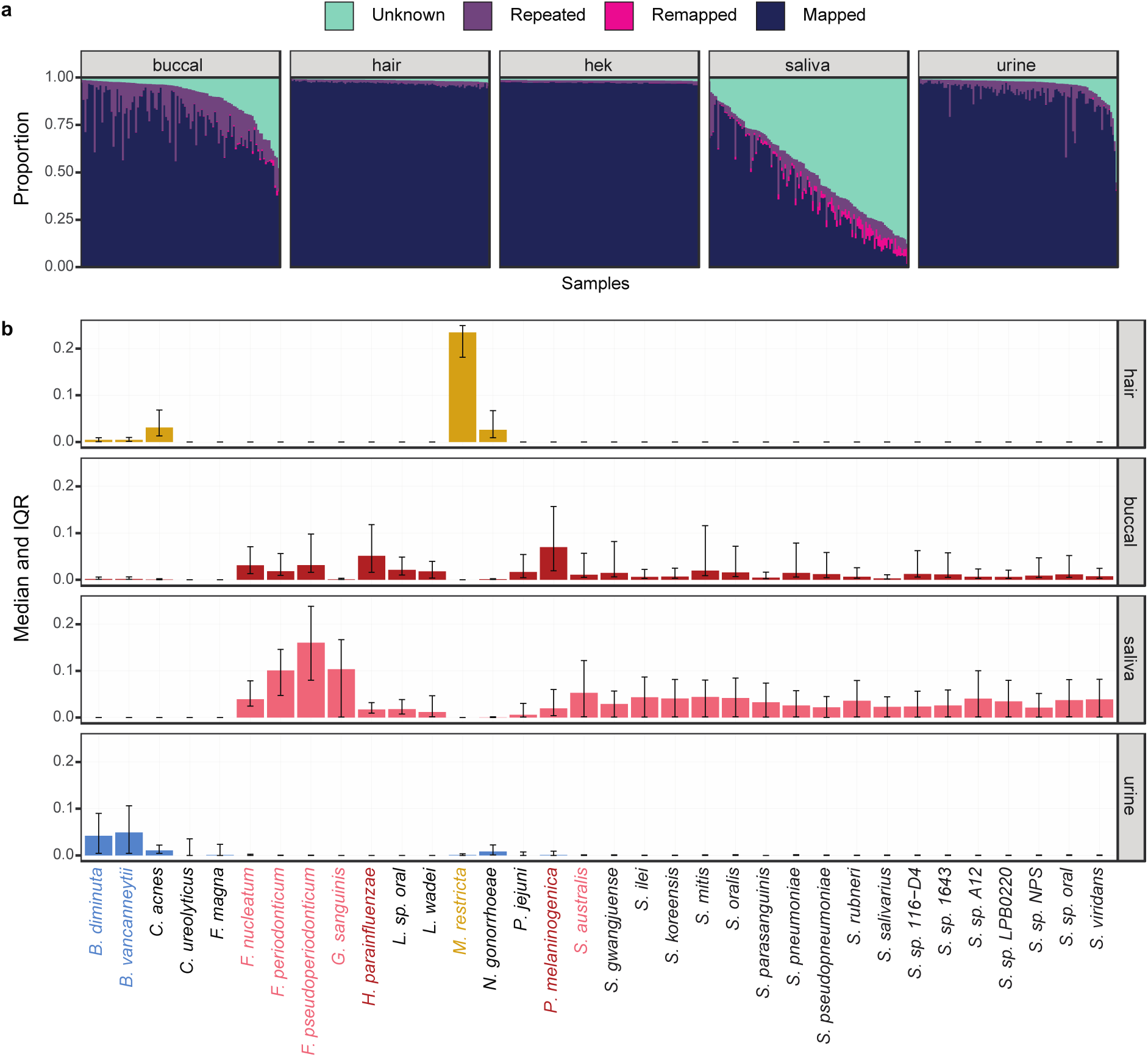
Classification of unmapped reads in noninvasive samples. **a.** Proportion of reads per sample. Mapped = aligned to hg38. Remapped = aligned to microbial species using Decontaminer. Repeated = highly abundant reads identified by FastQC. Unknown = reads not mapped, remapped, or highly abundant. **b.** Normalized proportion of reads remapping per species for each tissue. The top 0.5% most abundant microbes are shown. Highlighted species have a median abundance > 0.05 for that tissue.

### Sources of gene expression variance between individuals and noninvasive tissue types

Next, we investigated gene expression variance due to noninvasive tissue type or individual. Saliva samples largely failed to pass post-sequencing quality standards and were excluded from several analyses for this reason. Using a mixed linear model to delineate biological and technical contributors to gene expression variability, we found tissue type is the main driver of variance across samples (Figure 3a). These results were supported by PCA, which segregated samples primarily by tissue type, with hair forming a separate, distinct cluster from the other tissues (Figure 3b).

**Figure 3.**
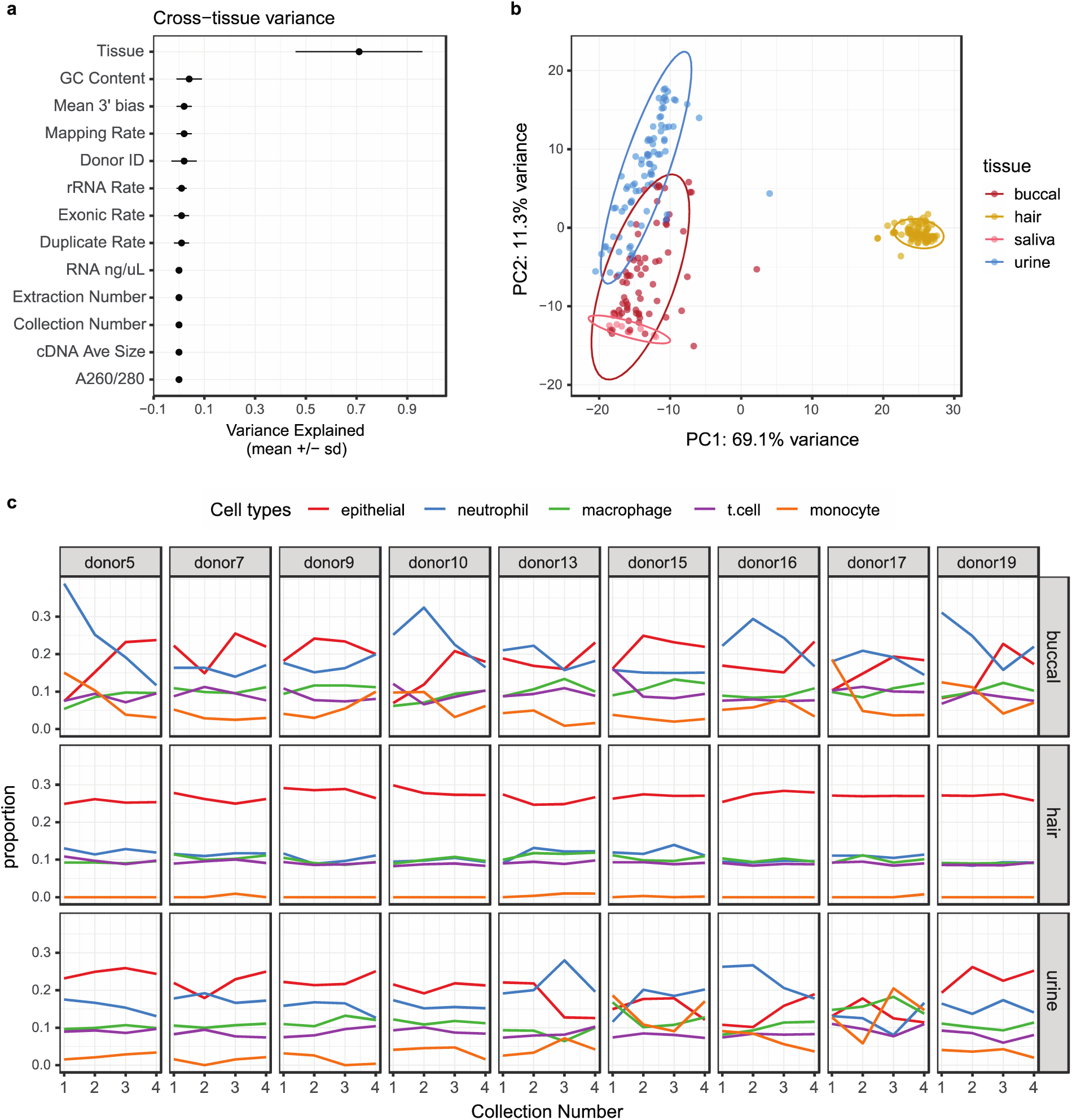
Technical and biological sources of variance in noninvasive samples. **a.** Factors contributing to variance in gene expression across tissues as determined by mixed linear modeling. **b.** Principal component analysis using DESeq2 normalized counts and the top 1000 most variable genes. **c.** GEDIT cell type proportion estimates across collections per donor. Only donors with samples passing QC for all collections are displayed here. Niche cell types were collapsed into larger categories (Supp. Fig. 7b), and the top 25% most abundant cell type categories across tissues are shown.

Because underlying cell type composition frequently explains gene expression variance across samples and tissues^15, 16^, we deconvolved our noninvasive samples using GEDIT^45^ and the provided BlueCodeV2 single cell reference. From this we observed hair is primarily composed of epithelial cell types, buccal and urine capture both epithelial and immune cells, and saliva mostly contains neutrophils and monocytes. Looking across tissues, donors, and collections, hair was highly consistent in cell type abundance estimates, both within and across donors (Figure 3c, Supp. Fig. 7a). On the other hand, urine and buccal samples were more variable. In these samples, cell type composition was occasionally consistent within and across donors, but there were also patterns where cell type composition changed across collections for the same donor (e.g. donor 5, buccal), showed a different pattern of abundance compared to other donors (e.g. donor 16, urine), or completely lacked any consistency (e.g. donor 17 urine). When we used linear mixed modeling to identify biological and technical sources of gene expression variance within tissues, the individual donor was the primary contributor in buccal and urine (Supp. Fig. 6). Both of these results indicate that cell type compositions sampled from urine and buccal samples are potentially highly variable and donor specific. For hair, the relative contribution of technical factors and the donor of origin to gene expression variance is similar. As previously described, hair follicle data quality matches gold-standard RNA-sequencing data, which together with the low variance in cell type composition leads to highly consistent gene expression profiles.

### Noninvasive tissue characteristics suggest potential invasive tissue type proxies

To investigate the biological similarity of noninvasive tissues to known, invasive sample types, we compared gene expression, splicing, and cell type enrichment patterns to GTEx. To do so, we first selected a single noninvasive sample per donor and per tissue with the highest protein-coding depth. Representative GTEx tissues were chosen based on k means clustering, and 19 samples of each tissue type were randomly selected. Both the noninvasive and GTEx samples were downsampled (see methods) to 5 million read counts to normalize for differences in total sequencing depth.

Using this data we projected the noninvasive samples onto the GTEx PCA space to observe global patterns of gene expression similarity (Figure 4a). Hair clusters closely with esophageal mucosa and skin, saliva is proximal to spleen, blood, and EBVs, and buccal and urine are intermediaries between these groups. We repeated this analysis using splicing events generated from rMATS^46^ (Figure 4b). From this we recapitulate similar clustering patterns observed for expression.

**Figure 4.**
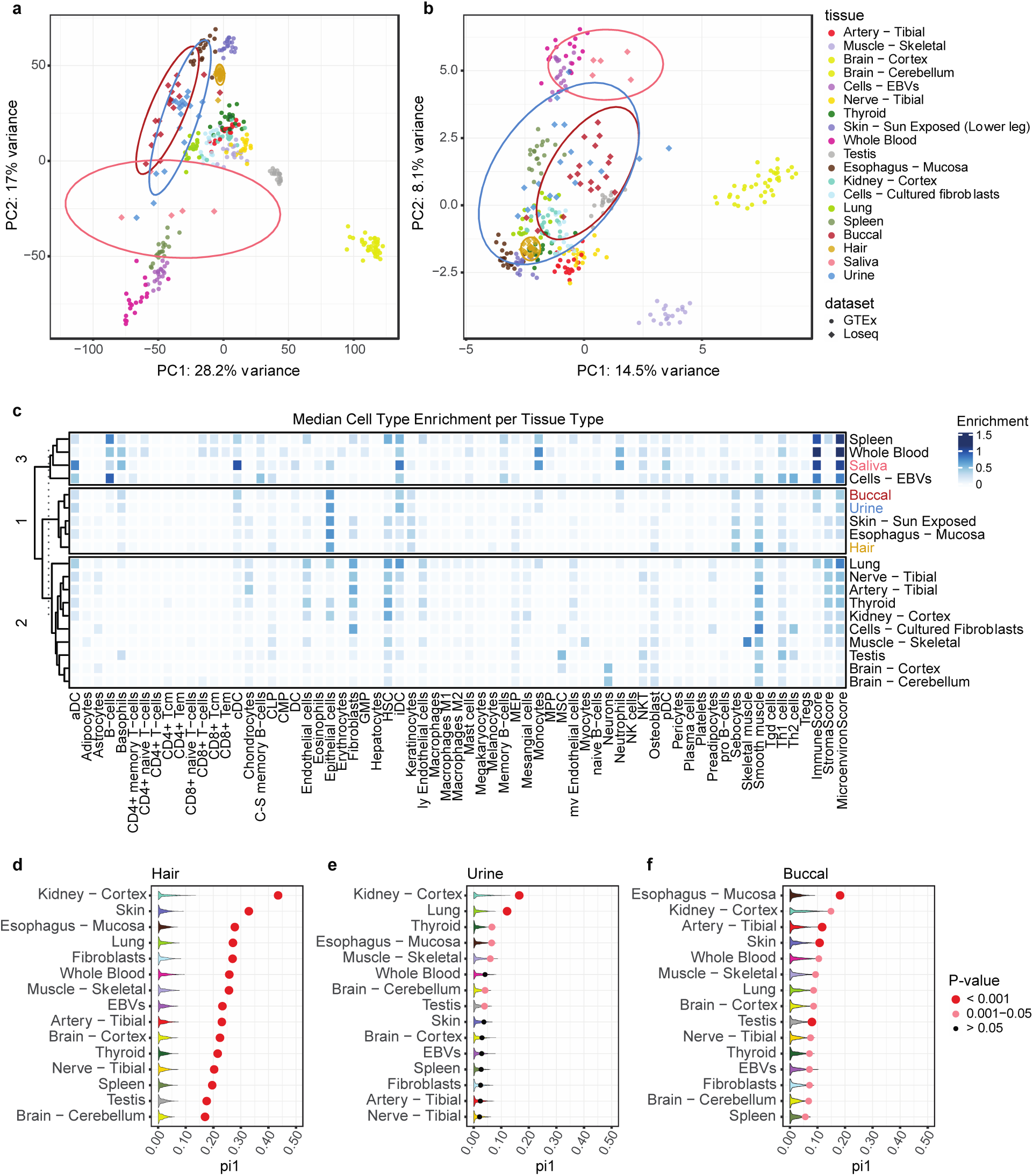
Comparison of noninvasive samples to the GTEx dataset. **a.** Noninvasive sample types projected onto the GTEx expression PCA space. Counts were normalized using DESeq2, centered and scaled, and the top 1000 most variable genes were used. Ellipses represent 95% confidence intervals. **b.** Noninvasive sample types projected onto the top 1000 most variable rMATS splicing events in GTEx. **c.** xCell cell type enrichment estimates per tissue. Tissues are clustered using k-means clustering. **d. e. f.** GTEx eQTL replication estimates for hair, urine, and buccal samples. Dots show π_1_ calculated by selecting significant GTEx gene-variant pairs from the noninvasive data with sizing indicating permutation p-value significance. Violin plots show null π1 distributions generated from allele-frequency matched, randomly selected gene-variant pairs. 1000 permutations were performed.

Since these clusters may reflect similarities in underlying cell types, we investigated this question by using xCell^47^ to calculate cell type enrichment scores. Here we used xCell because it is the most comprehensive cell type database available, thus enabling analysis of diverse tissues and biospecimens, and we observed high concordance in cell type estimates between GEDIT and xCell for cell types present in both references (Supp. Fig. 8). From this analysis we replicated the same tissue clustering we observed by PCA except using cell type enrichment estimates (Figure 4c). Because cell type sharing is highly predictive of shared gene regulatory mechanisms^26–29^, this suggests gene expression regulatory mechanisms present in invasive tissues may be captured noninvasively.

To further explore how noninvasive samples may capture genetic regulatory variants discovered in postmortem tissues, we assessed replication of GTEx eQTLs in buccal, hair, and urine samples. With our sample size being insufficient for full eQTL discovery, we looked for enrichment of low p-values in our noninvasive dataset for the eVariant-eGene pairs previously discovered in GTEx. A null distribution was generated by randomly sampling allele-frequency matched eVariant-eGene pairs from our noninvasive data, and we calculated π_1_, an estimate of the true positive rate, in both the null datasets and when selecting the significant GTEx eVariant-eGene pairs. In hair, we find significant replication of GTEx pairs across all studied GTEx tissues, with kidney cortex and skin showing the highest degree of replication (π_1_ = 0.44 and π_1_ = 0.33, respectively, Figure 4d). Buccal and urine are less homogenous tissue types, thus further decreasing our power especially as our sample size does not allow highly efficient approaches to correct for latent variation^48^. As such, they showed less clear signal across all tissues (Supp. Fig. 9). However, we did observe most significant enrichment for kidney cortex eVariant-eGene pairs in urine and esophageal mucosa signal enrichment in buccal (π_1_ = 0.17 and π_1_ = 0.18, respectively, Figures 4e and 4f). Of note, kidney cortex has a low sample size relative to other tissues in GTEx and thus little power for discovery of more subtle eQTL effects. Thus, it is unclear whether the high replication of kidney eQTL signal across noninvasive tissues is due to similar biology or an abundance of common and/or high effect size eQTLs in this tissue. In all, our results suggest noninvasive tissues capture cell types and gene expression regulatory mechanisms present in invasive tissue types and may provide insight into disease processes affecting these tissues.

### Sex-specific differences in gene expression in noninvasive samples

Because biological and environmental contexts play a major role in expression regulation, we aimed to explore whether RNA-sequencing from noninvasive tissues may be used for this purpose. To this end, we tested for sex-based differential expression and the replication and biological role of the discoveries. Using edgeR^49^ and limma-voom^50^, we were able to identify 25 and 1032 sex-based differentially expressed genes in hair and urine, respectively (Supp. Fig. 10a). In comparing hair to sun-exposed skin, 8 of the 25 significant hits were previously seen in GTEx and are highlighted in Figure 5a. Looking across all GTEx tissues, 17 of the 25 genes were previously observed (Supp. Fig. 10b). Running FGSEA^51^ on the hair results showed significant enrichment for E2F targets and G2M checkpoint pathways in females (Figure 5b), and these are central to regulating the cell cycle and proliferation^52, 53^. Though nonsignificant, Wnt signaling, the top hit for males, has previously been reported to play a key role in hair loss prevention^54^. In urine, 33 of the 1032 significant findings were seen in kidney cortex, and, overall, 582 were differentially expressed in any GTEx tissue (Figure 5c, Supp. Fig. 10b). Notably, estrogen response is greatly enriched in females (Figure 5d). Estrogen signaling is central to many physiological processes in the kidney and is considered potentially protective against many renal diseases, though much remains unknown^55^. This analysis demonstrates the potential for noninvasive samples to elucidate underlying biology in a variety of potential contexts and assays.

**Figure 5.**
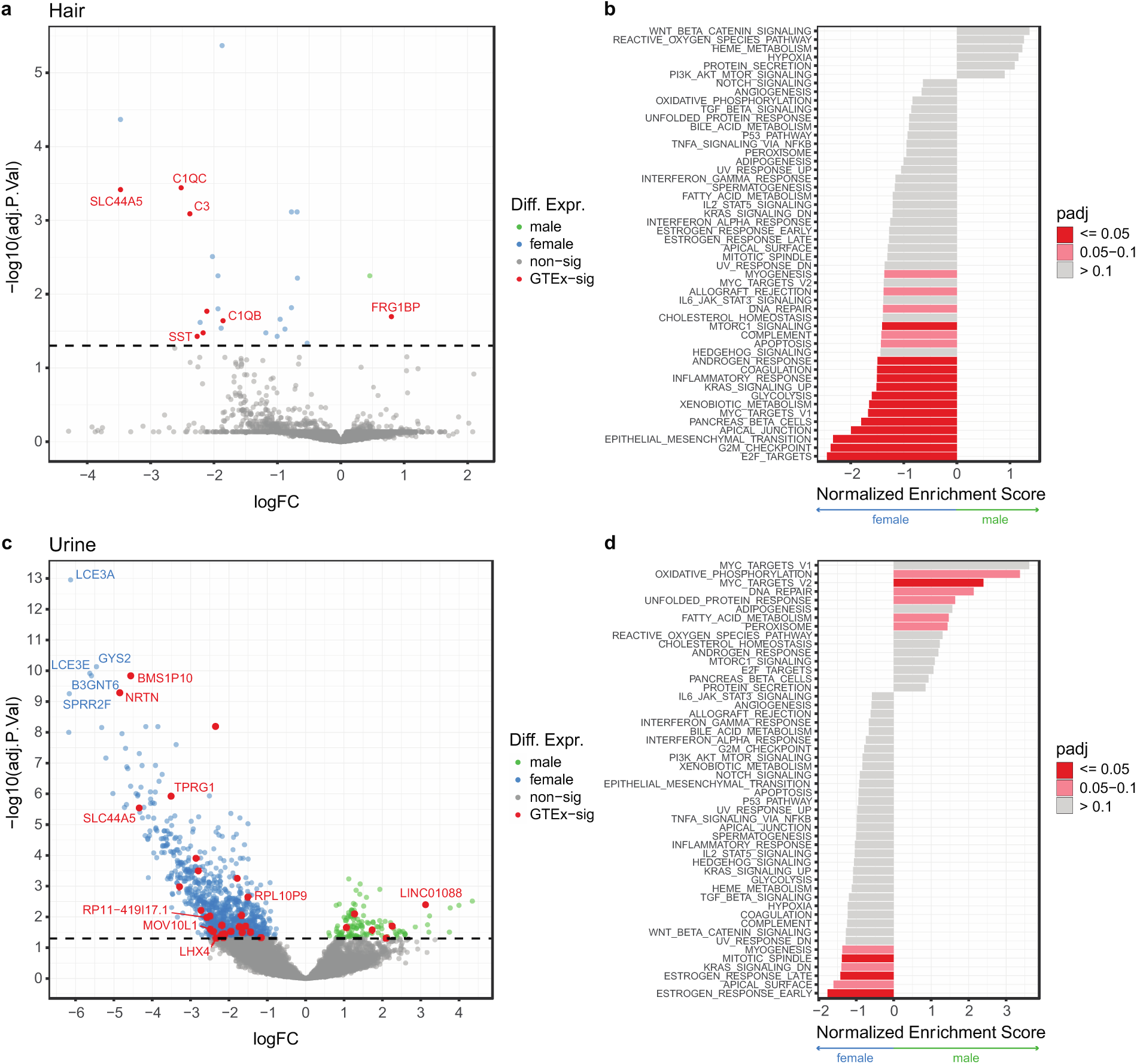
Sex-based expression differences in noninvasive samples. **a.** Volcano plot of sex-based differentially expressed genes in hair. Genes highlighted in red are replicated sun-exposed skin findings in GTEx. Dotted line indicates 0.05 significance threshold. **b.** Hair FGSEA of all genes ranked by z-score and using the Hallmark Gene set from MSigDB. **c.** Sex-based differentially expressed genes in urine cell pellets. Genes highlighted in red are replicated kidney cortex findings in GTEx. **d.** Urine FGSEA of all genes ranked by z-score and using the Hallmark Gene set from MSigDB.

### Noninvasive samples may be leveraged for disease-relevant applications

Allele specific expression (ASE) analysis compares allelic expression levels within the same individual, and it is an important tool for investigating rare and cis-regulatory variation, nonsense mediated decay, and genomic imprinting^56^. Here we quantified ASE using heterozygous sites across tissues. From this, we observe anticipated reference allele ratio patterns per individual (Supp. Fig. 11a) and depending on the SNP annotation (Supp. Fig. 11b), and we show robust nonsense mediated decay for stop-gain variants across all collections for buccal, hair, and urine samples (Figure 6a). This suggests noninvasive sampling may be used to identify gene-disrupting variants that are a common focus of genetic diagnosis in rare disease.

**Figure 6.**
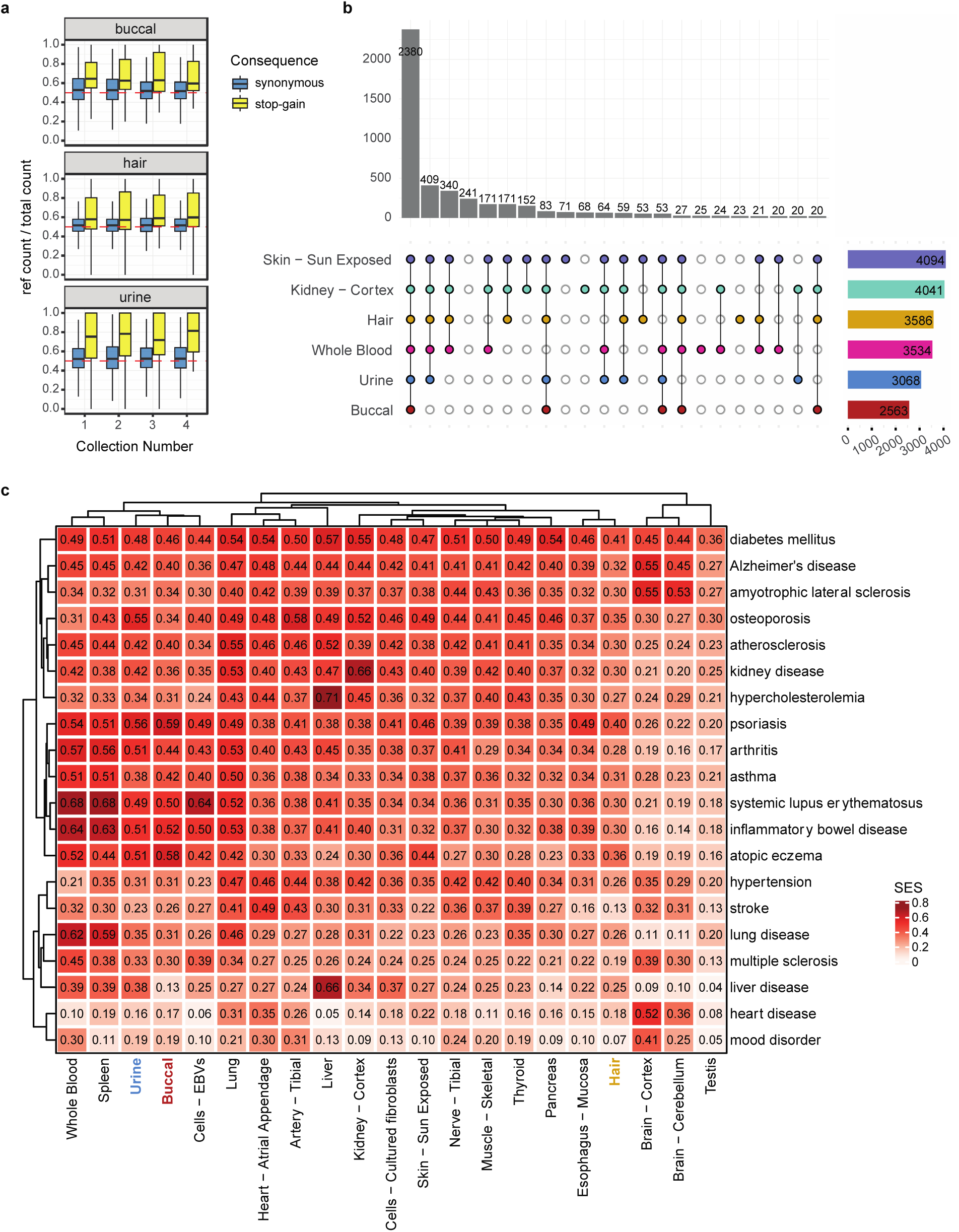
Use of noninvasive samples in disease-relevant analyses. **a.** ASE for annotated stop-gain variants vs synonymous. Only sites with > 16 total counts were included. **b.** Genes with median expression > 0.1 TPM in a tissue were intersected with the OMIM gene set. Shown is the intersection of OMIM genes captured across tissues. **c.** Capture of common disease signals in the OpenTargets database. SES = Σ(evidence scores of disease genes expressed in a tissue)/ Σ(evidence scores of disease genes expressed in any included tissue).

Application of noninvasive samples to rare and common disease was evaluated using the OMIM^57^ and OpenTargets^58^ repositories. For Mendelian disease, our samples captured a median of 80% (hair), 70% (urine), and 55% (buccal) of genes above a 0.1 TPM threshold (Supp. Fig. 13a). This capture is consistently high for hair samples across collections but shows donor-dependent consistency for urine (Supp. Fig. 13b). Clustering samples by median OMIM gene expression with GTEx recapitulated our prior proxy observations (Supp. Fig. 13c). In Figure 6b, we show the overlapping gene sets for Mendelian genes with a median expression greater than 0.1 TPMs in a given tissue after employing GTEx expression thresholds within that tissue. The selected GTEx tissues shown are relatively minimally invasive or identified as most similar by clustering and eQTL replication. We see the vast majority of genes are captured in both noninvasive and invasive tissue types, indicating noninvasive samples may be a suitable biospecimen for studying gene expression and regulatory processes in many rare disease applications. Notably, we observe a subset of genes expressed in noninvasive samples that are not captured in whole blood (ex. 171 in hair, skin, and kidney only; 83 in all tissues except for blood). This suggests diseases where the primary tissue type affected is more akin to noninvasive samples may benefit more from the use of noninvasive sampling versus whole blood collection. Importantly, recent work from others investigating clinically accessible tissues further supports the conclusions we draw here^59^. In all, efforts to improve clinical genomics studies using transcriptomics may be further augmented by use of noninvasive samples, especially where invasive surgical sampling of tissues primarily affected is often not possible.

Looking at common disease, we first selected the most general OpenTargets ontology category for every disease included. We filtered for genes with greater than 5 sources of evidence and with tissue elevated specificity from the Human Protein Atlas database^60^ (Supp. Fig. 12). Disease enrichment was calculated by summing together OpenTargets gene evidence scores (summed evidence score, SES) for genes with median expression greater than zero in a given tissue and dividing by the total possible summed evidence score. From this, we found buccal and urine captured diseases with a strong immunological component, much like whole blood and spleen (Figure 6c). Urine additionally showed strong signal for kidney disease (SES = 0.42). Hair performed best for skin-related diseases (psoriasis SES = 0.40), but overall did not show strong enrichment for any particular disease. These results suggest noninvasive samples bear promise for use in disease-relevant studies while providing the advantage of study designs with potentially longitudinal monitoring and greater enrollment.

## DISCUSSION

Discovery from transcriptomic data and its use in precision medicine is considerably limited by cost and access to biologically applicable biospecimens^18, 61^. As a result, most transcriptome studies have lagged behind GWAS in sample size, which now often include hundreds of thousands of individuals, and studies still need to increase enrollment from non-European populations. Further, disentangling causation and finding context-specific disease mechanisms is challenging using a single collection time point^17, 62^. To address these limitations, we sought to investigate low-cost, noninvasive RNA-sequencing as an alternative approach. From our study we observed hair follicles and urine cell pellets provide the highest quality data and perform best in functional genomics and clinical applications.

A primary advantage of noninvasive biospecimens over blood-related specimens to the transcriptomics field is the set of cell types captured. Shared cell type composition corresponds with shared regulation of gene expression and splicing^15, 16, 21, 22^, and a major limitation of blood-related samples is that they represent a highly tissue-specific set of cell types. In noninvasive samples we observed greater similarity by expression, splicing, and genetic regulation to invasive GTEx tissues relative to GTEx whole blood. Cell type deconvolution analysis estimated noninvasive tissues to contain epithelial cells, myocytes, stromal cells, and others, all of which are unable to be captured using blood and play a key role in mechanisms of many diseases.

Several considerations should be taken into account when deciding to use noninvasive tissues. Generally, ease of sample collection and consistency of library preparation performance and quality is biospecimen-dependent. Additionally, the feasibility of clinical use and longitudinal study design varies depending on the tissue type. We observed high failure rates using buccal swabs and saliva, and though the data yielded from samples passing quality standards provides valuable insights and the samples themselves are simplest to collect, we believe use of these biospecimens should be reserved for specialized applications where health status of the oral cavity and/or upper gastrointestinal tract is primarily under study.

Though we found urine cell pellets to yield high quality RNA, the pellet itself may be minimal and difficult to visualize for some donors. This is especially true for healthy donors, who tend to shed fewer cells into their urine^29^, and could introduce bias into future study designs if special care is not taken when using this tissue. Similarly, we found urine cell pellet RNA quantity to be variable and donor dependent. Others have aforementioned single cell approaches for urine specimens^30, 31^, and we anticipate further development of these methods will better control for sampling inconsistency. Here, we showed urine cell pellets capture genetic regulatory mechanisms seen in the kidney as well as gene expression signatures relevant to kidney disease and diseases mediated via kidney functions. Given the enormous health burden kidney disease poses in the US and worldwide, and the central role the kidney plays in many diseases^63^, methods for noninvasively monitoring kidney function and enabling early diagnosis could meaningfully improve morbidity and mortality. In addition to the analyses performed here, others have proposed laboratory protocols for propagating cells collected from urine for use in identifying disease mechanisms and novel treatments, or, in the case of stem cells, developing autologous cell therapies^32–34^. Overall, given further optimization, we expect urine holds the greatest potential for clinical use and discovery.

Hair follicles perform robustly using any library preparation and exhibit low technical variance across collections and donors. It should be noted that hair follicle collection does require additional training of personnel not necessarily needed for the other biospecimens. Also, fine versus coarse hair type played a role in determining the ease of collection, and we do observe slight differences in yield depending on the donor, though this did not impact sample performance. We do foresee the need to explore collection of hair follicles from other parts of the body when head hair is not available, and it is likely necessary in future studies to collect additional information regarding the use of cosmetics and medications applied to the head and skin. Here, we showed hair follicles result in consistent quality, cell types, and expression profiles across collections, and, despite our low sample size, we found significant replication of previously observed eQTLs across all GTEx tissues. Together, these findings suggest hair is a highly robust biospecimen with potentially broad applications, and, because of its consistency, biological perturbations due to disease, treatment, or other environmental exposures will likely be observable in clinical and longitudinal settings.

Notably, across all noninvasive tissues we observed a large majority of Mendelian disease genes were expressed. The ease and decreased invasiveness of noninvasive proxy biospecimens could facilitate greater use of transcriptome analyses in diagnosing rare, genetic disease^59^, however, further work is needed to explore this possibility.

Our findings will need to be validated using larger, more diverse, and clinical cohorts. We expect noninvasive tissues may reduce Eurocentric sampling bias and enable sampling from more vulnerable populations, but this expectation will need to be measured against future study enrollment. Additionally, we used bulk RNA-sequencing, and the performance and features of noninvasive samples using single cell methods will need to be optimized and evaluated.

## CONCLUSION

In this study we aimed to establish whether noninvasive sampling may be used to scale transcriptomic studies. From our work, we were able to characterize many of the technical and biological features of four possible noninvasive samples, and we showed their potential utility in both transcriptomic and disease-related applications. Overall, we find hair follicles and urine cell pellets to be the most promising biospecimens, and we propose advantages in terms of cost and study designs for pursuing noninvasive sampling. In all, noninvasive RNA-sequencing offers meaningful improvements to current transcriptomic approaches that could enable dramatic scaling in sample size and increased discovery potential. This scaling would bring closer parity with GWAS via transformational increases in power, thus better positioning transcriptomic studies for use in diagnostic and clinical applications.

## METHODS

### Noninvasive sample collection

IRB approval was obtained for the study. 19 participants were recruited and consented, and four total collections of four tissue types were completed. The first collection occurred 6 months prior to confirm piloted procedures were ready to scale, and the remaining 3 collections were performed within a 2-4 week window per participant. 75 samples of each tissue type were obtained (1 participant provided only 3 collections). A detailed noninvasive sample collection protocol is provided on protocol.io (DOI: dx.doi.org/10.17504/protocols.io.kqdg3pjzzl25/v1).

### Library preparation and sequencing

RNA extraction procedures are unique to each tissue type, and thus were performed separately for each tissue. Collections were randomized across extractions. All samples were prepared using our in-house method with 15 samples of each tissue type in duplicate. Takara Bio SMART-seq v4 and Illumina stranded mRNA prep kits were prepared according to the manual and using a random subset of 12 and 16 samples of each tissue type, respectively. In total, 6 library preparation plates were prepared, and 2 HEK cell samples were included in triplicate on each library prep plate. This resulted in 508 total samples. 485 samples passed yield and size pre-sequencing quality parameters (>2nM yield and <600 bp average size). Samples were pooled by tissue type with HEK cell samples randomized across the library pools. Libraries were sequenced 2×150 bp on a Novaseq 6000 S4 flow cell. A detailed sample preparation protocol for RNA-extraction and our in-house method is provided on protocol.io (DOI: dx.doi.org/10.17504/protocols.io.kqdg3pjzzl25/v1).

### Alignment, quantification of gene counts, and quality assessment

Adapter sequences were removed using Trimmomatic 0.36^64^. Sequences were aligned to build 38 of the human genome with Gencode v35 annotation using STAR 2.7.3a^65^ and Samtools 1.9^66^ set to GTEx mapping parameters^18^. Marking of duplicate reads was done using Picard 2.23.7^67^, and gene counts were quantified using featureCounts from Subread 1.6.5^68^. QC results output from FastQC 0.11.3^69^, STAR, and RNA-SeQC 2.3.6^39^ were consolidated using MultiQC 1.8^70^. QC filtering based on standard sequencing quality metrics or based on protein coding and lncRNA depth thresholds were found to be largely redundant (Supp. Fig. 2), and thus depth of mapped reads was used for its simplicity. Genes were determined to be expressed in a tissue and included in downstream analyses if raw counts >= 8 and TPMs >= 0.1 in >= 20% of samples within a given tissue type.

### Unmapped reads analysis

Unmapped reads were output to fastq files during alignment. These reads were remapped using DecontaMiner 1.4^40^. 23,488 bacterial, 21 fungal, and 11,120 viral genome references were downloaded as suggested in the Installation and User Guide. Default parameters were used to remove low quality, human ribosomal and mitochondrial reads. For BLASTn alignments to the reference databases, bacterial and fungi parameters included minimum length = 50bp and gaps and mismatches = 2bp. Gaps and mismatches were increased to 5bp for viral genome remapping. Organisms were left unfiltered during initial remapping settings (Supp. Fig. 5a). For the analysis, we normalized the results by genomic length of the remapped species and by number of remapped reads per sample. We selected the top 0.5% of remapped species across all tissue types.

Technical and biological sources of unmapped reads were investigated by first removing all Decontaminer remapped reads from the unmapped fastqs, and then using FastQC to identify overrepresented sequences in the remaining reads. These sequences were compiled across all tissues and samples into a list of 707 sequences. Command line tools were used to filter and quantify the overrepresented read counts per sample. Computational and manual comparison to primer and adapter sequences as well as comparisons across preps and tissues were used to delineate the potential sources of the reads (Supp. Fig. 5d).

### Downsampling

For analyses involving downsampling, binomial sampling was performed 5 times on the raw, unfiltered counts matrix and then taking the average. Binomial probability of success was set to the (desired depth)/(original depth), number of observations set to the total gene number, and trials set to the gene counts per a given gene. Samples were QCed for protein coding and lncRNA depth prior to downsampling. For comparisons across different library preparations, all samples passing a threshold depth of 1 million reads mapped to the human genome were included and then subsequently downsampled to 1 million (Supp. Fig. 2a,b,c). Loseq analyses were thresholded and downsampled to 2.5 million protein coding depth (Supp. Fig. 2d,e,f), with the exception of the cell type deconvolution analysis which was thresholded and downsampled to 5 million. All GTEx comparisons were made with both Loseq and GTEx thresholded and downsampled to 5 million.

### Cross-preparation comparison

Median TPMs per gene were calculated within a given tissue, prep, and replicate group, and the Pearson correlation across replicate groups 1 and 2 for a given tissue and prep was quantified (Supp. Fig. 3). Overlap of gene expression capture was evaluated by taking the median TPMs within a tissue and prep, filtering genes with zero median expression, and determining the gene overlap using ComplexUpset 1.3.1^71, 72^ in R 3.6.0 (Supp. Fig. 3). Principal component analysis was performed using DESeq2 1.26.0^73^ VST normalized counts and by selecting for the top 500 most variable genes. Variance attributable to tissue and prep was done by performing linear regression per PC (PC ∼ tissue + prep) followed by ANOVA with p-value correction based on the number of PCs tested (Supp. Fig. 4).

### Loseq cross-sample variance assessment

Principle components analysis was done across tissues using DESeq2 1.26.0 VST normalized counts and by selecting for the top 1000 most variable genes. Correlation of technical variables with PCs was investigated via linear regression. VariancePartition 1.21.6^74^ was used to identify sources of gene expression variance within and across tissue types. The cross tissue VariancePartition model included collection, extraction, donor id, and tissue as random variables and rRNA rate, mapping rate, duplicate rate, exonic rate, 3’ bias, RNA concentration, a260/280, cDNA size, and GC content as fixed variables. Within tissue models were the same except tissue type was dropped as a variable (Supp. Fig. 6).

### Cell type deconvolution of noninvasive samples

Deconvolution was done using GEDIT 1.7^45^ and the provided BlueCodeV1.0.tsv reference matrix. For the final analyses, cell types were collapsed into broader umbrella categories by adding the estimated proportions together (Supp. Fig. 7b). Only the top 25% most abundant cell types per tissue were considered when looking across collections, and only donors with all 4 collections passing QC in urine, hair, and buccal were included in the final plot. All donors are plotted in the supplement (Supp. Fig. 7a).

### PCA projection of noninvasive samples onto GTEx

For Loseq, the sample with the highest protein coding and lncRNA depth passing the 2.5 million threshold was selected per donor and per tissue type (19 samples per hair and urine, 17 buccal, 5 saliva), and both Loseq and GTEx were downsampled to 5 million. 19 samples of each representative (see xCell section) GTEx tissue were randomly sampled. Counts were VST normalized using DESeq2 1.26.0. Principal components analysis was run on centered and scaled GTEx counts using the top 1000 most variable genes. The resulting PCA loadings were multiplied by GTEx and Loseq centered and scaled counts, and this data was plotted as shown in the main figure.

The splicing PCA was performed in the same manner except using an exon inclusion level matrix generated by rMATS 4.1.2^46^ as input. Splicing events with zero inclusion for any sample in a tissue were excluded. The rMATS results were filtered for events found to be significantly different across the select GTEx tissues and with inclusion levels greater than 2 standard deviations beyond the average inclusion (0.3). Again, PCA was run for the top 1000 most variable events in GTEx, and the loadings were multiplied by the GTEx and Loseq centered and scaled inclusion levels.

### Cell type enrichment analysis of noninvasive and GTEx samples

75 samples were sampled from each GTEx tissue to approximately match the number of Loseq samples included (75 hair, 63 urine, 25 buccal, 5 saliva). GTEx and Loseq TPMs were deconvolved for enrichment using 64 cell type signatures in xCell 1.1.0^47^ in R 3.6.0. Select GTEx tissues were chosen using gene expression clustering of median TPMs per GTEx tissue. GTEx groups were established using k means, and the tissue with the highest sample size per group was selected as representative. Kidney cortex, esophageal mucosa and lung were added based on their proximity to noninvasive tissues we studied. Clustering of cell type enrichment was done by taking the median enrichment score per tissue and then using k means in ComplexHeatmap 2.2.0^75^.

### GTEx eQTL replication analysis

Participants provided their genotyping data from 23andme (9 donors), Ancestry (2 donors), and Gencove (8 donors) platforms. Array data was imputed using the 1000 genomes phase 3 reference^76^ and the Sanger Imputation Service^77^. Eagle 2.4.1^78^ and the 1000 genomes reference were used for VCF phasing. Monomorphic alleles, alleles with MAF < 0.05, multiallelic sites and indels were excluded from all analyses. This imputed, phased, and filtered VCF was used for eQTL and ASE analyses.

For the genotyping PCA, LD pruning was performed with PLINK 1.90-b3.29^79^ with a window size of 50, shift of 5, and r squared cutoff of 0.2. Only SNPs with a 100% genotyping rate and HWE 1e-5 were included. Ancestry was imputed by merging the donor VCF with the 1000 genomes VCF, excluding any individuals with relatedness >= 0.0625, running smartpca with eigensoft 6.1.3^80^, and using k-nearest neighbors to infer the ancestry population of our donors. Running smartpca on the donor samples alone revealed genotyping PC1 corresponded with ancestry, while PC2 corresponded with genotyping panel. These were included as covariates in the eQTL analysis, as well as expression PCs with variance explained > 15% that accounted for global changes in gene expression (PC1 for hair and buccal and PCs 1 and 2 for urine). Counts were TMM and inverse normalized and filtered for genes with raw counts >= 6 and TPMs >= 0.1 (as is the GTEx standard). eQTL mapping was done on a per tissue basis using TensorQTL v.1.0.5 with the window set to 1MB (following the GTEx parameters).

To calculate replication, each GTEx tissue was filtered for the top eVariant-eGene pair per gene with MAF >= 0.05, qvalue <=0.05, and with effect size greater than the minimum observed in the lowest powered GTEx tissue (kidney cortex = 0.32). This was intersected with the pairs discovered in the noninvasive dataset, and pi1 was calculated using qvalue 2.18.0^81^ and R 3.6.0. Null datasets were of the same size as the overlapping significant GTEx pairs set and were generated by sampling allele-frequency matched eVariant-eGene pairs from the noninvasive data 1000 times. Pi1 was calculated per each dataset. Significance for enrichment was determined based on permutation p-value calculations (Supp. Fig. 9).

### Differential expression analysis and FGSEA

Sex-based differential expression analysis was performed on a per tissue basis using edgeR 3.28.1^49^ and Limma-Voom 3.42.2^50^. Counts were filtered based on GTEx parameters and TMM normalized prior to analysis. GTEx sex-based differential expression results^82^ were retrieved from the GTEx Portal (https://gtexportal.org/home/datasets) for overlap comparisons (Supp. Fig. 10b). For noninvasive tissues, a named list of sex-based differentially expressed genes and their t-scores was input into FGSEA 1.12.0^51^. The Hallmark gene sets file was obtained from Molecular Signatures Database (http://www.gsea-msigdb.org/gsea/msigdb/collections.jsp#H)^83^ and used for the analysis.

### Loss of function detection using ASE

Allele-specific expression was calculated using ASEReadCounter 4.0.1.1^56^ and using the imputed, phased, and filtered VCF described in the GTEx replication analyses. Sites with fewer than 16 total counts were filtered from the analysis. The reference counts divided by total counts was assessed across tissues and donors, and one donor was removed due to extreme ratios and thus potential genotyping errors (Supp. Fig. 11a). Ensembl Variant Effect Predictor 5.28.1^84^ was used to annotate variant consequences, and the ratio of reference to total allele counts was compared given these annotations (Supp. Fig. 11b).

### OMIM Mendelian disease gene overlap

Genes with Mendelian inheritance were downloaded from the OMIM database^57^ (https://www.omim.org/). Tissues were filtered were genes meeting minimum GTEx expression thresholds, and then the remaining gene set was overlapped with OMIM genes (Supp. Fig. 13). ComplexUpset 1.3.1 was used to compare genes captured across noninvasive and select GTEx tissues.

### OpenTargets evaluation of disease-relevancy

Data was retrieved from the OpenTargets database^58^. Disease ids were selected by choosing the broadest ontological category specific to a given disease (as provided on the OpenTargets platform). For the analysis, the association file incorporating all sources of evidence was used, and disease genes were included only if there were 5 or more sources of evidence (Supp. Fig. 12a). Loseq samples were thresholded and downsampled to a protein coding and lncRNA depth of 5 million, and samples with the highest depth per donor and tissue were selected. GTEx was similarly downsampled and 19 samples of each GTEx tissue were randomly selected. GTEx tissues were chosen based on their relevance to selected diseases. Genes were filtered based on GTEx parameters, and the median TPMs per tissue was calculated. 10,986 tissue-elevated genes were obtained from the Human Protein Atlas^60^ (https://www.proteinatlas.org/humanproteome/tissue/tissue+specific), and each tissue was additionally filtered for tissue-elevated genes. In the end, the top 3,411 most expressed, tissue-elevated genes present in a tissue were analyzed, based on the tissue with the lowest number of genes passing post-expression and HPA filtering (Supp. Fig. 12b).

To calculate the summed evidence score, first, the original target overlap was calculated by intersecting the genes present across all tissues with the disease gene targets. Summing the evidence score for these genes resulted in the total possible score for a given disease. Then, the genes expressed in a particular tissue were overlapped with disease gene targets. The evidence scores for tissue-specific overlapping genes were summed together and then divided by the total possible score for a disease. This normalized score is the summed evidence score reported in the analysis (Supp. Fig. 12c).

## ACKNOWLEDGEMENTS

We would like to thank the members of the Lappalainen laboratory, most notably Kristina Buschur and Paul Hoffman, as well as William Mauck (Satija Laboratory, NYGC) and Stephanie Hao (Innovation Laboratory, NYGC). We thank the 19 donors for their participation and contribution to science.

## AUTHOR CONTRIBUTIONS

M.M., S.E.C, and T.L. designed the study. S.E.C. performed donor recruitment and enrollment. S.E.C. and M.M. optimized the in-house library preparation protocol. M.M. collected and prepared all donor samples for sequencing and performed or led all analyses herein. A.G. assisted in data generation. R.G. did the unmapped reads and splicing analyses. S.K. provided code, expertise, and supervision of eQTL analyses. M.M. wrote the manuscript. S.E.C., S.K., R.G., and T.L. edited the manuscript. All authors read and approved the final manuscript.

## FUNDING

M.M. was supported by the NHGRI (F30HG011194). S.K., and T.L. were supported by the NHLBI (R01HL142028). R.G., A.G., and T.L. were supported by the NIA (R01AG057422). S.E.C. was supported by the NHGRI (1K99HG009916).

## AVAILABILITY OF DATA AND MATERIALS

The data set generated and analyzed in the study is available on dbGaP (accession number posted upon publication). Noninvasive sample collection and in-house library preparation methods are provided on protocols.io (dx.doi.org/10.17504/protocols.io.kqdg3pjzzl25/v1). Analysis code has been deposited in the GitHub repository: https://github.com/LappalainenLab/loseq

## ETHICS, CONSENT, AND PERMISSIONS

IRB approval was obtained via both Columbia University and BRANY for the study, and all 19 participants herein consented to participate.

## COMPETING INTERESTS

T.L. advises and has equity in Variant Bio, and advises Goldfinch Bio. and GSK. S.E.C. is a cofounder, chief technology officer, and stock owner at Variant Bio.

## Notes

### Summary of Updates

Abstract, introduction, and title revised to clarify impact of the work. Additional quantitative results provided in the library preparation results section.

