## Supplemental Materials for "Evaluation of noninvasive biospecimens for transcriptome studies"

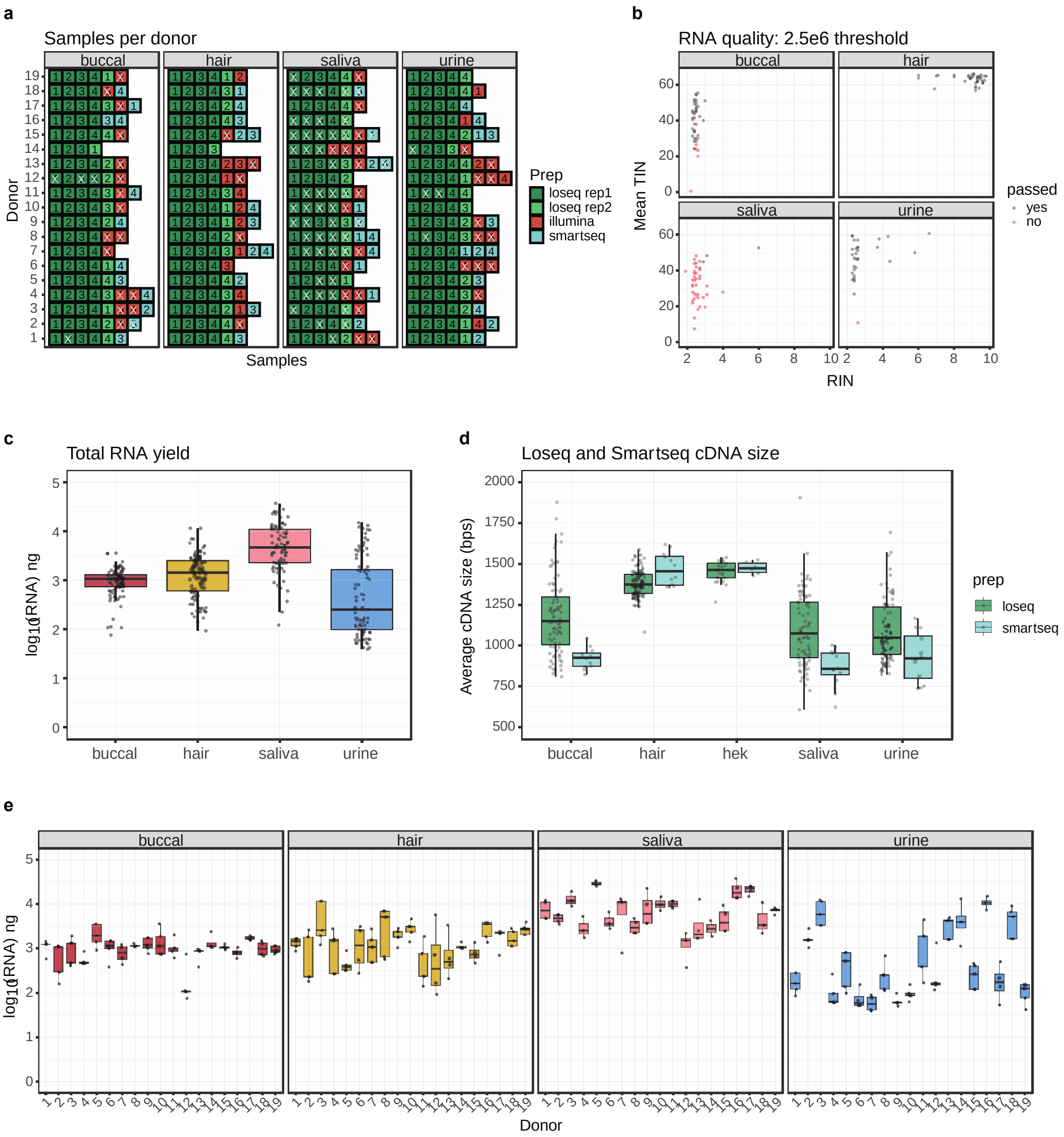
a
Samples per donor
buccal
hair
saliva
urine
Donor
Samples
Prep
loseq rep1
loseq rep2
illumina
smartseq
b
RNA quality: 2.5e6 threshold
buccal
hair
saliva
urine
Mean TIN
RIN
passed
yes
no
c
Total RNA yield
log10(RNA) ng
buccal
hair
saliva
urine
d
Loseq and Smartseq cDNA size
Average cDNA size (bps)
buccal
hair
hek
saliva
urine
prep
loseq
smartseq
e
buccal
hair
saliva
urine
log10(RNA) ng
Donor

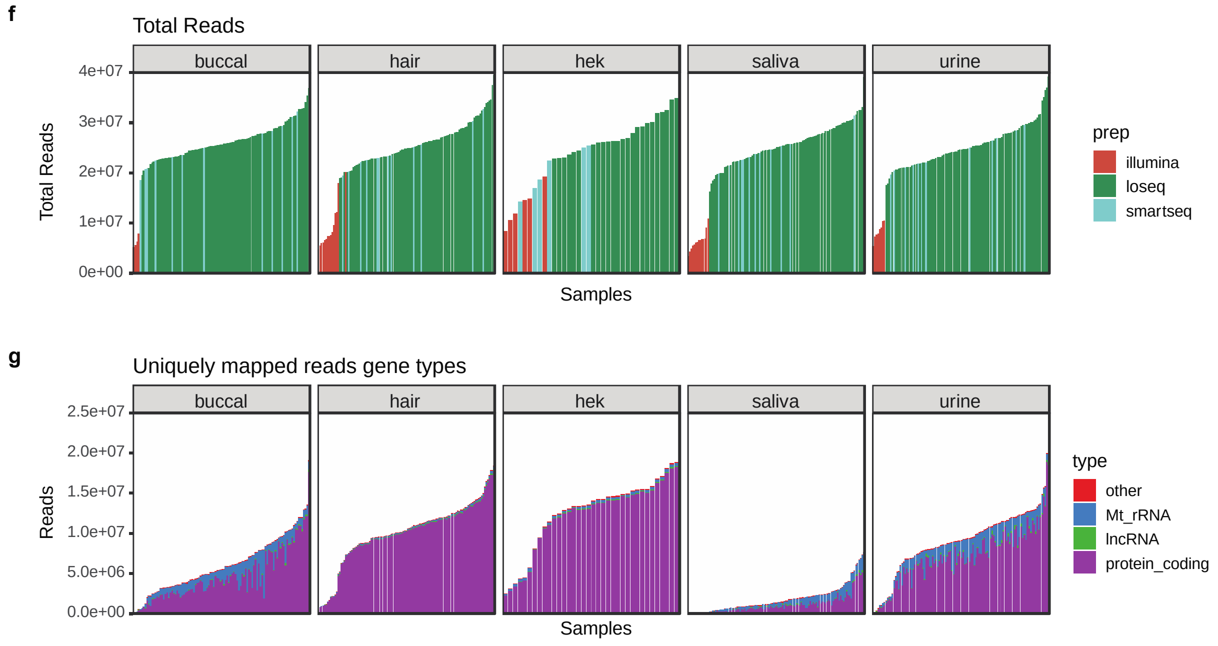


**Supplementary Figure 1. a.** Outcome of library preparation QC per donor, collection, and preparation. The crosses indicate a failed sample, and the numbers correspond to the collection. **b.** Measured RIN vs computationally-derived transcript integrity number (TIN) per sample. Passing is determined by 1 million protein-coding depth threshold. **c.** RNA yield distribution per tissue type. Buccal median = 1054.2ng, Q1 = 735.7ng, Q3 = 1292.55ng. Hair median = 1444.1ng, Q1 = 606.2ng, Q3 = 2530.5ng. Saliva median = 4705.4ng, Q1 = 2278.5ng, Q3 = 10920.35ng. Urine median = 253.4ng, Q1 = 98ng, Q3 = 1642.2ng. **d.** cDNA average size for Loseq and SmartSeq preparations across noninvasive tissues. **e.** RNA yield per donor and tissue. Each data point is a collection. Buccal Levene P = 0.05, Hair Levene P = 0.01, Saliva Levene P = 0.2, Urine Levene P = 0.0002. **f.** Total reads sequenced per sample, colored by prep. **g.** Categorization of gene types for uniquely mapped reads mapping to genes across tissues.

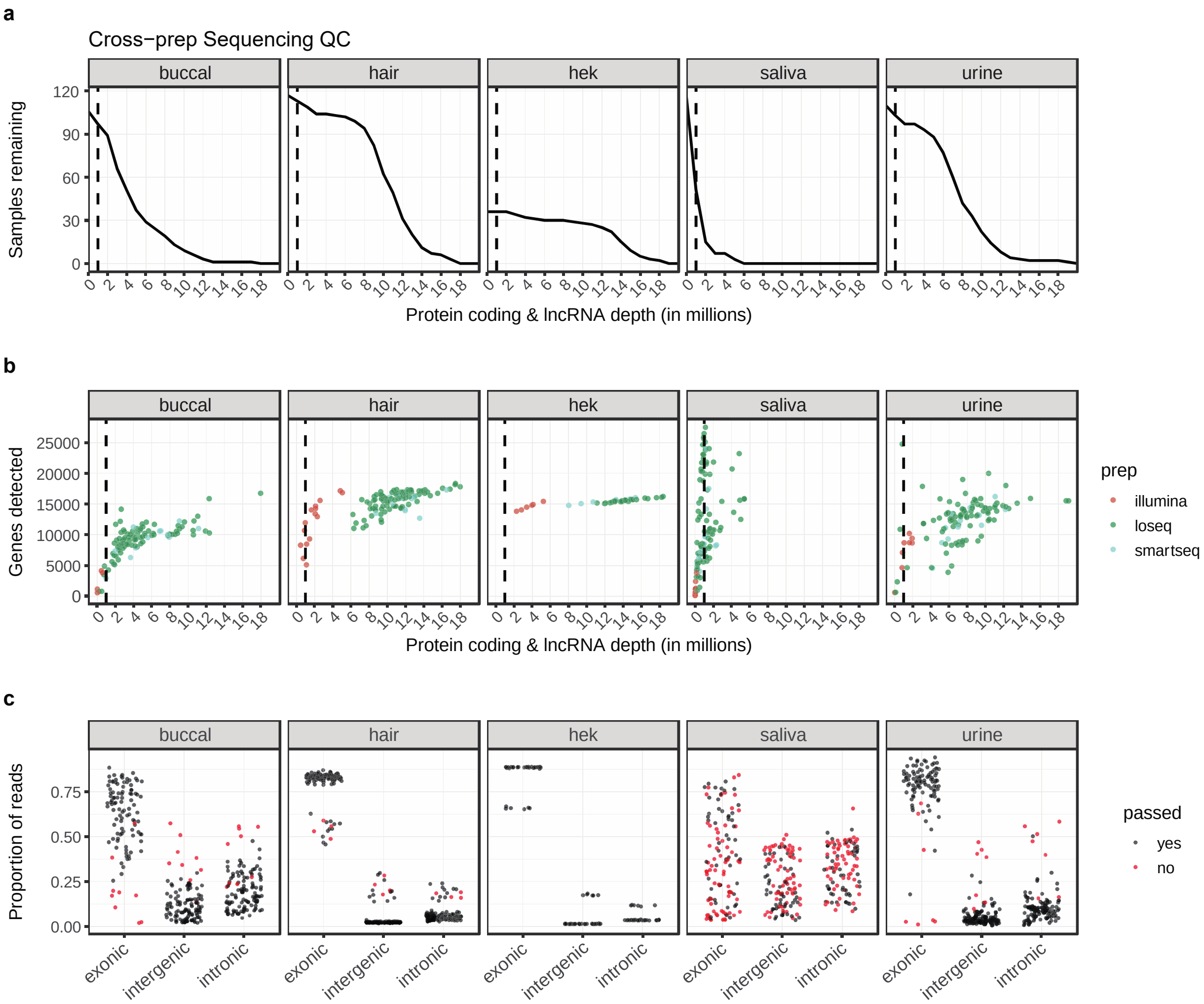
a
Cross−prep Sequencing QC
buccal
hair
hek
saliva
urine
Samples remaining
Protein coding & lncRNA depth (in millions)
b
Genes detected
prep
illumina
loseq
smartseq
c
Proportion of reads
exonic
intergenic
intronic
passed
yes
no

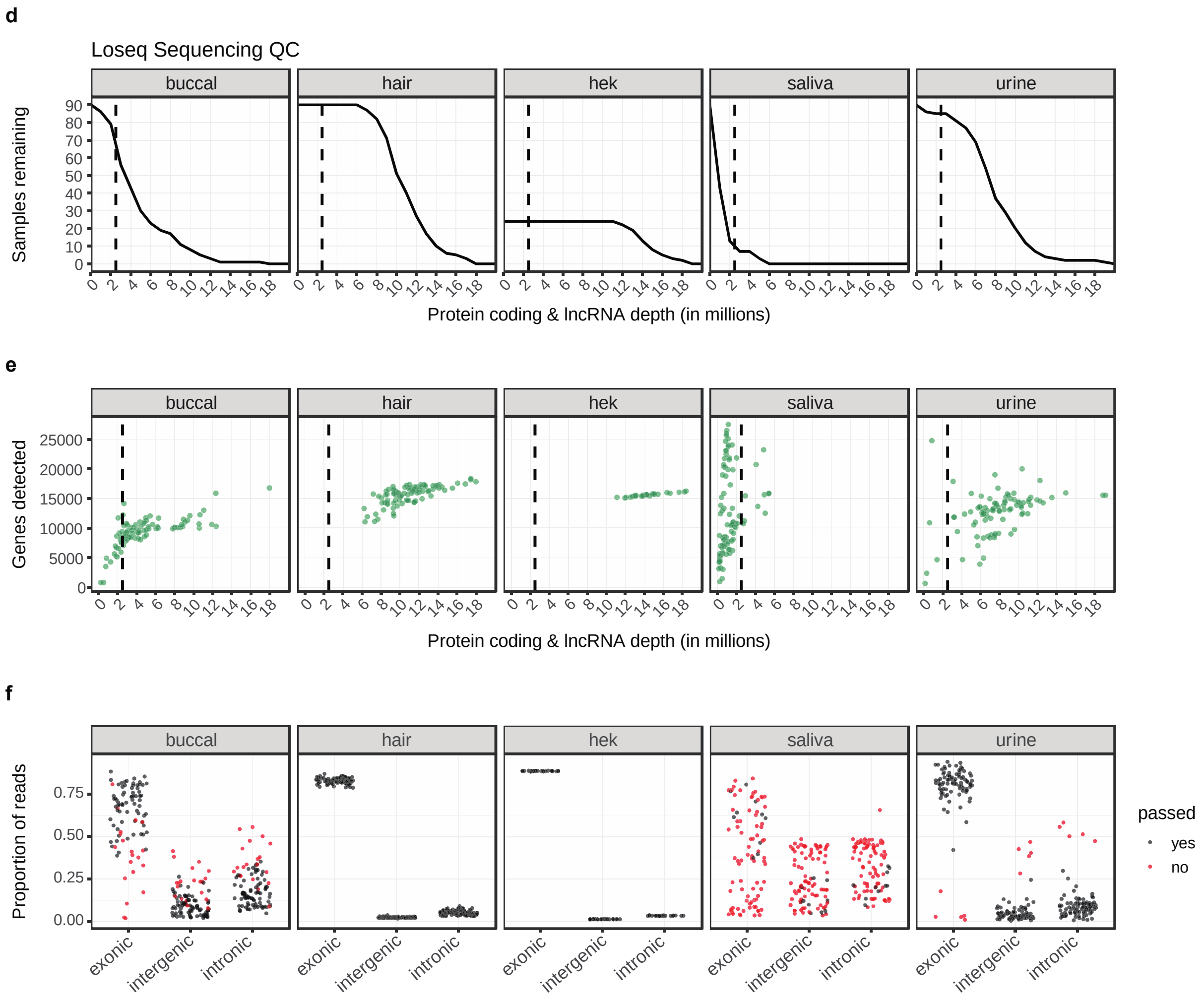


**Supplementary Figure 2. a-c** quality statistics for all samples across all library preparations: **a. b.** Samples remaining and genes detected depending on depth threshold. Dotted line indicates preparation QC threshold of 1 million. **c.** Proportion of reads mapping to exonic, intergenic, and intronic genomic features. Pass indicator is based on preparation QC threshold of 1 million. **d-f** quality statistics for only Loseq samples: **d. e.** Samples remaining and genes detected depending on depth threshold. Dotted line indicates Loseq QC threshold of 2.5 million. **f.** Proportion of reads mapping to exonic, intergenic, and intronic genomic features. Pass indicator is based on Loseq QC threshold.

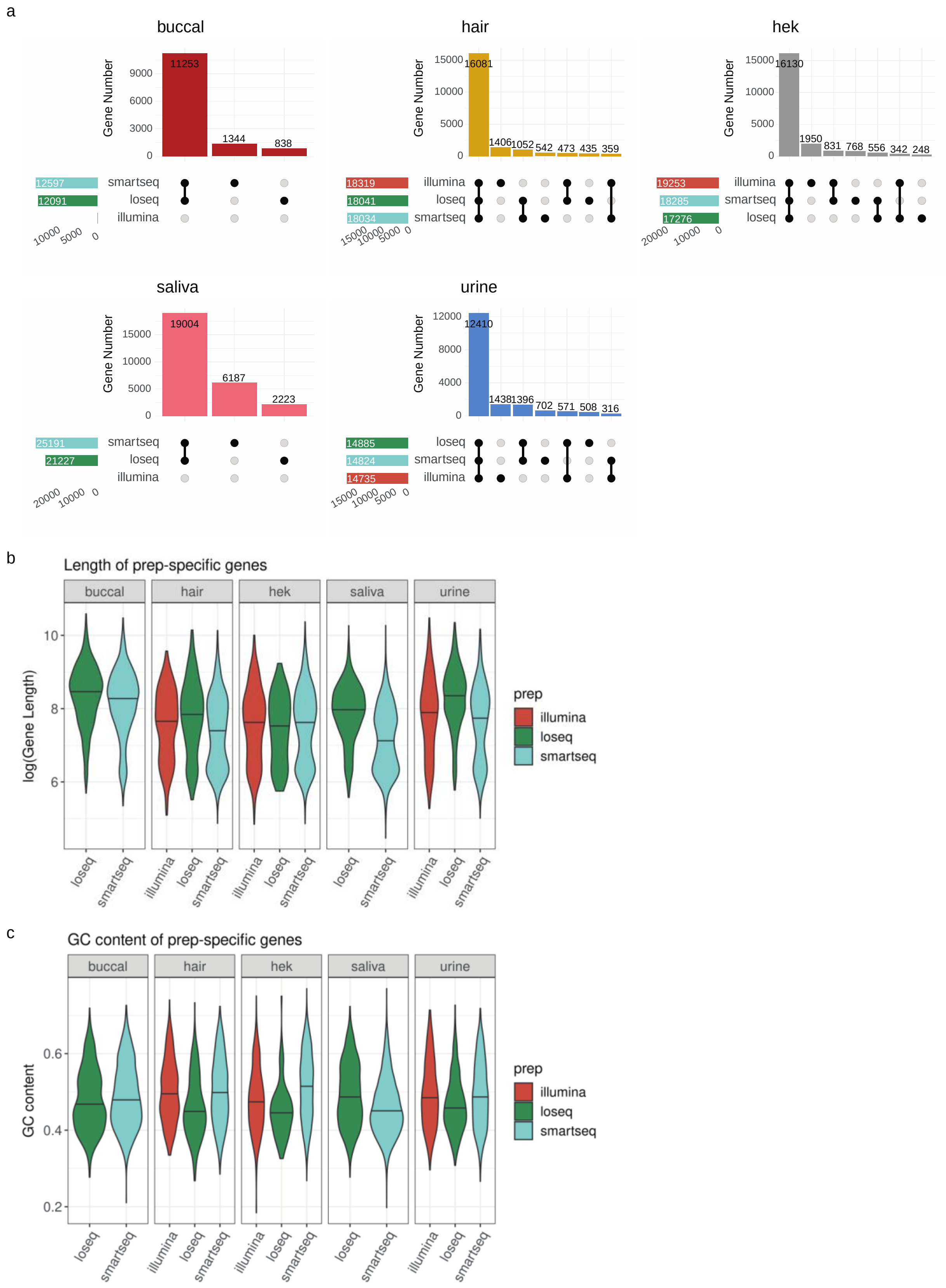
a
buccal
Gene Number
11253
1344
838
smartseq
loseq
illumina
12597
12091
hair
16081
1406
1052
542
473
435
359
18319
18041
18034
hek
16130
1950
831
768
556
342
248
19253
18285
17276
saliva
19004
6187
2223
25191
21227
urine
12410
1438
1396
702
571
508
316
14885
14824
14735
b
Length of prep-specific genes
log(Gene Length)
prep
illumina
loseq
smartseq
c
GC content of prep-specific genes
GC content

d

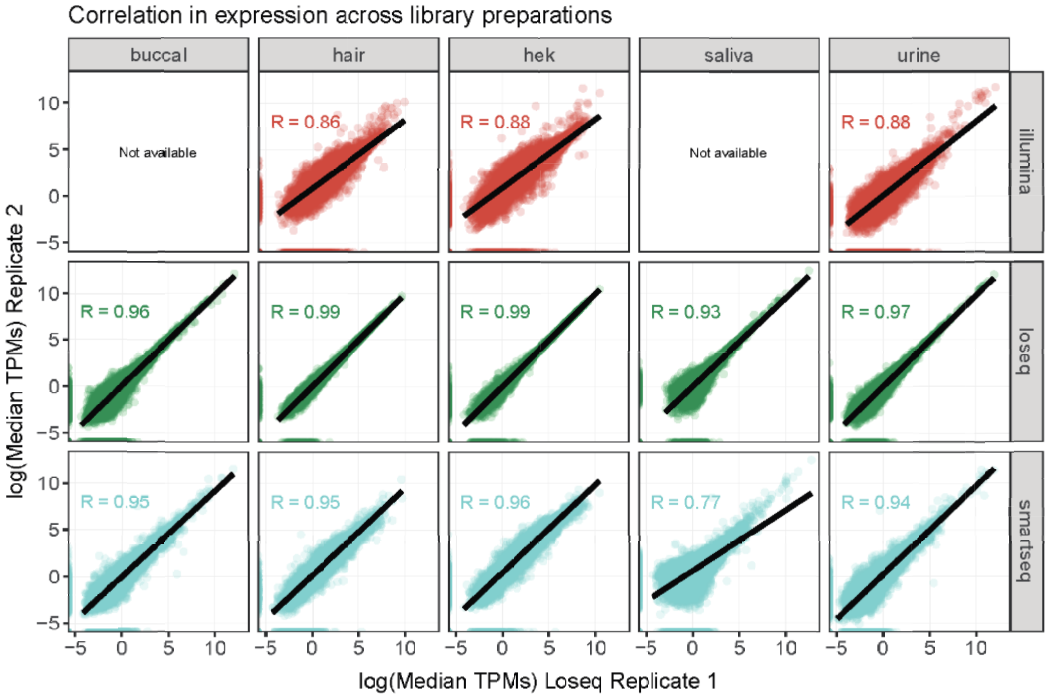


**Supplementary Figure 3. a.** Intersection of genes with greater than zero median expression across library preparations. **b. c.** Comparison of gene length and GC content of genes uniquely captured in a given preparation. **d.** Comparison of median gene expression levels of library preparation replicates. The median of all replicate 1 samples (15) is compared against the median of all replicate 2 samples (15) per library preparation. The spearman correlation between these is shown.

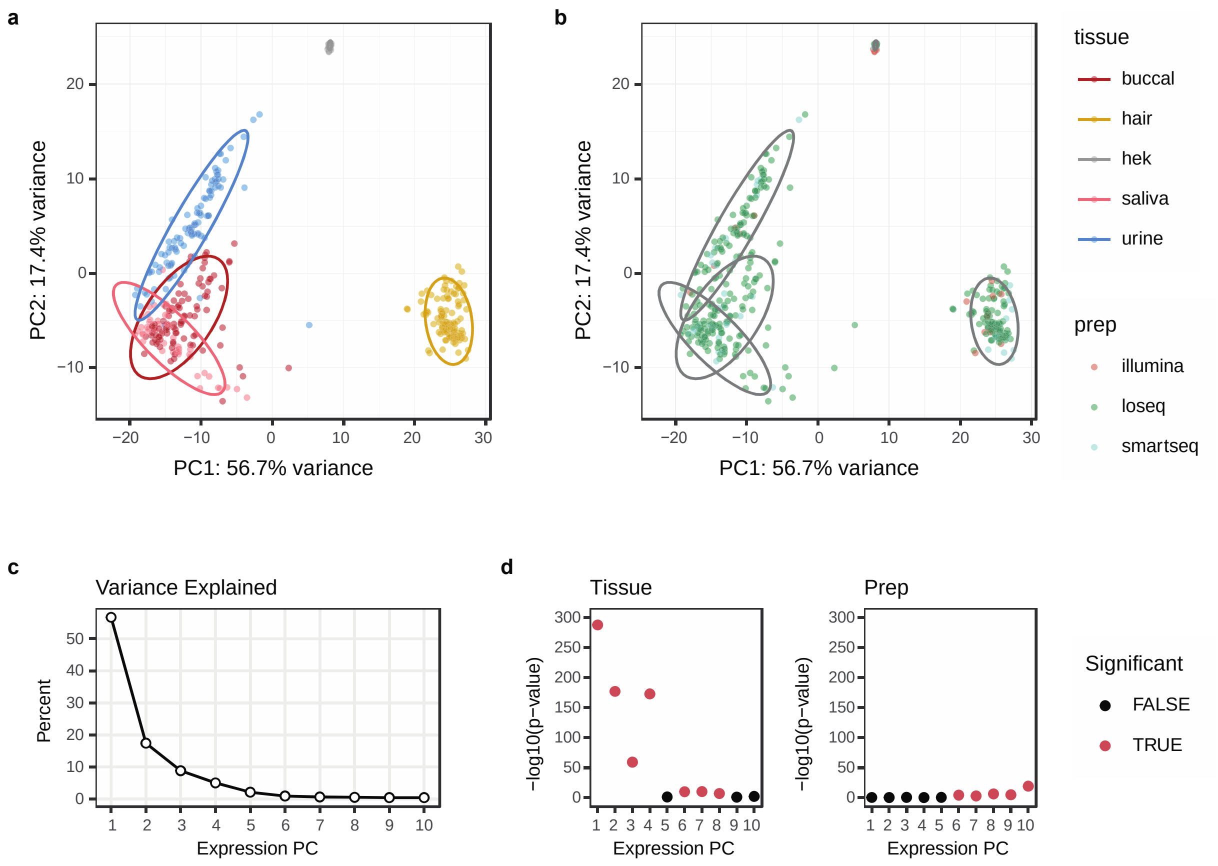


**Supplementary Figure 4. a. b.** Principal Component Analysis of all samples passing preparation QC thresholds. **c.** Percent variance explained per PC. **d.** ANOVA results for PC ~ tissue and PC ~ preparation. P-values are Bonferroni-corrected for the number of PCs tested (10).

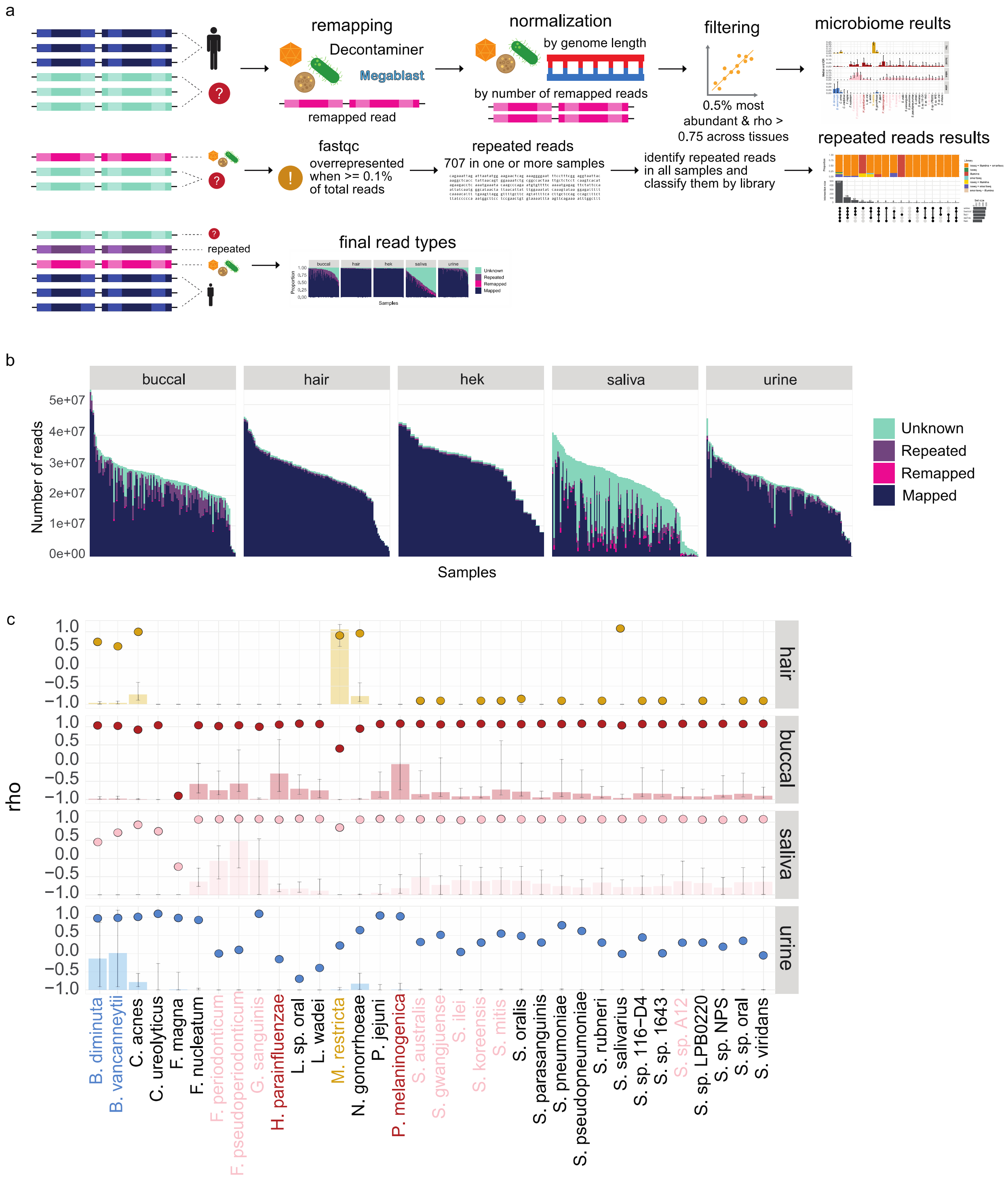
a
remapping
Decontaminer
Megablast
remapped read
normalization
by genome length
by number of remapped reads
filtering
0.5% most abundant & rho > 0.75 across tissues
microbiome reults
repeated reads results
fastqc
overrepresented when >= 0.1% of total reads
repeated reads
707 in one or more samples
identify repeated reads in all samples and classify them by library
repeated
final read types
b
buccal
hair
hek
saliva
urine
Number of reads
5e+07
4e+07
3e+07
2e+07
1e+07
0e+00
Samples
Unknown
Repeated
Remapped
Mapped
c
rho
1.0
0.5
0.0
−0.5
−1.0
hair
buccal
saliva
urine
B. diminuta
B. vancanneytii
C. acnes
C. ureolyticus
F. magna
F. nucleatum
F. periodonticum
F. pseudoperiodonticum
G. sanguinis
H. parainfluenzae
L. sp. oral
L. wadei
M. restricta
N. gonorrhoeae
P. jejuni
P. melaninogenica
S. australis
S. gwangjuense
S. ilei
S. koreensis
S. mitis
S. oralis
S. parasanguinis
S. pneumoniae
S. pseudopneumoniae
S. rubneri
S. salivarius
S. sp. 116−D4
S. sp. 1643
S. sp. A12
S. sp. LPB0220
S. sp. NPS
S. sp. oral
S. viridans

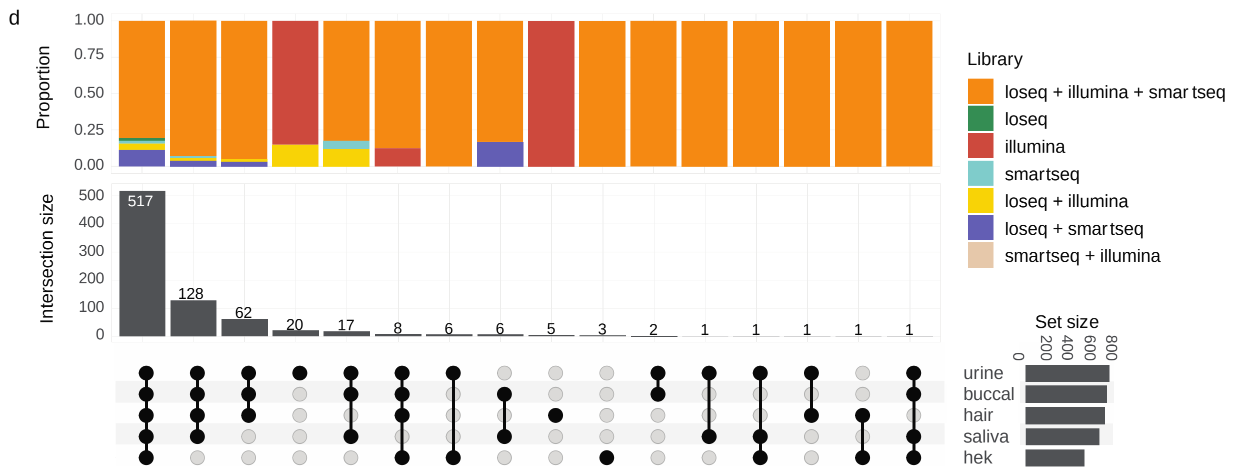


**Supplementary Figure 5. a.** Schematic of pipeline for assigning unmapped reads. Briefly, unmapped reads were remapped using decontaminer, normalized using a procedure akin to TPM normalization, then species of interest were filtered based on their abundance and replication across technical replicates. Remaining unmapped reads were investigated using FastQC. **b.** Total number of reads assigned to each category. Mapped = aligned to hg38. Remapped = aligned to microbial species using Decontaminer. Repeated = highly abundant reads identified by FastQC. Unknown = reads not mapped, remapped, or highly abundant. **c.** For each species included in the final analysis, spearman rank correlation between technical replicates is shown with the dots. Bar plot of species abundance with error bars is shown in the background for direct comparison (y-scale of abundance in Figure 2b). **d.** Breakdown of repeated sequence sharing across tissues and library preparations.

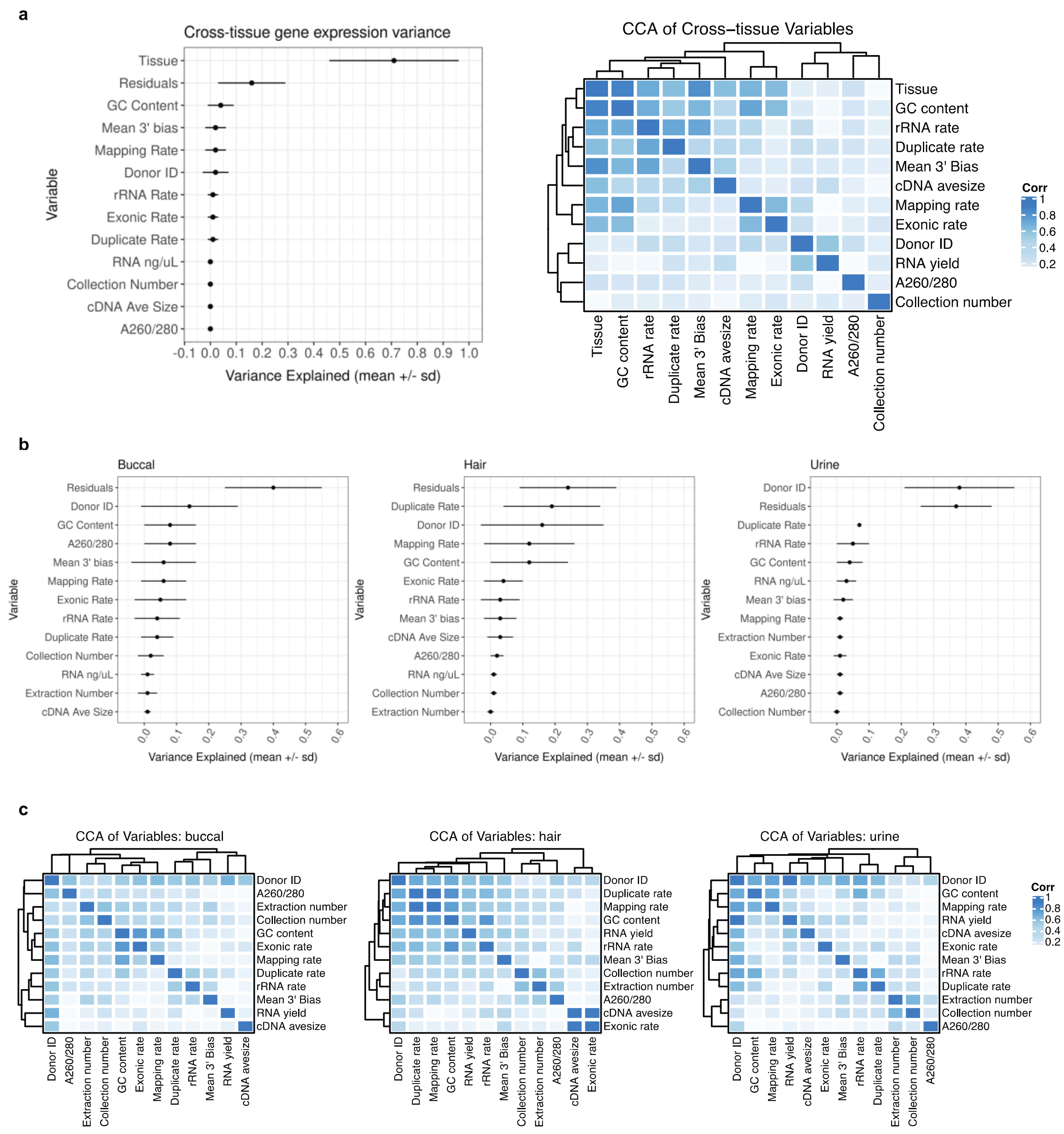


**Supplementary Figure 6. a.** Variance in gene expression across tissues explained by technical and biological variables. Canonical correlation analysis shows correlation between variables used. **b. c.** Variance in gene expression within each tissue explained by technical and biological variables. Canonical correlation analysis shows correlation between variables used. Only Loseq samples were included in these analyses.

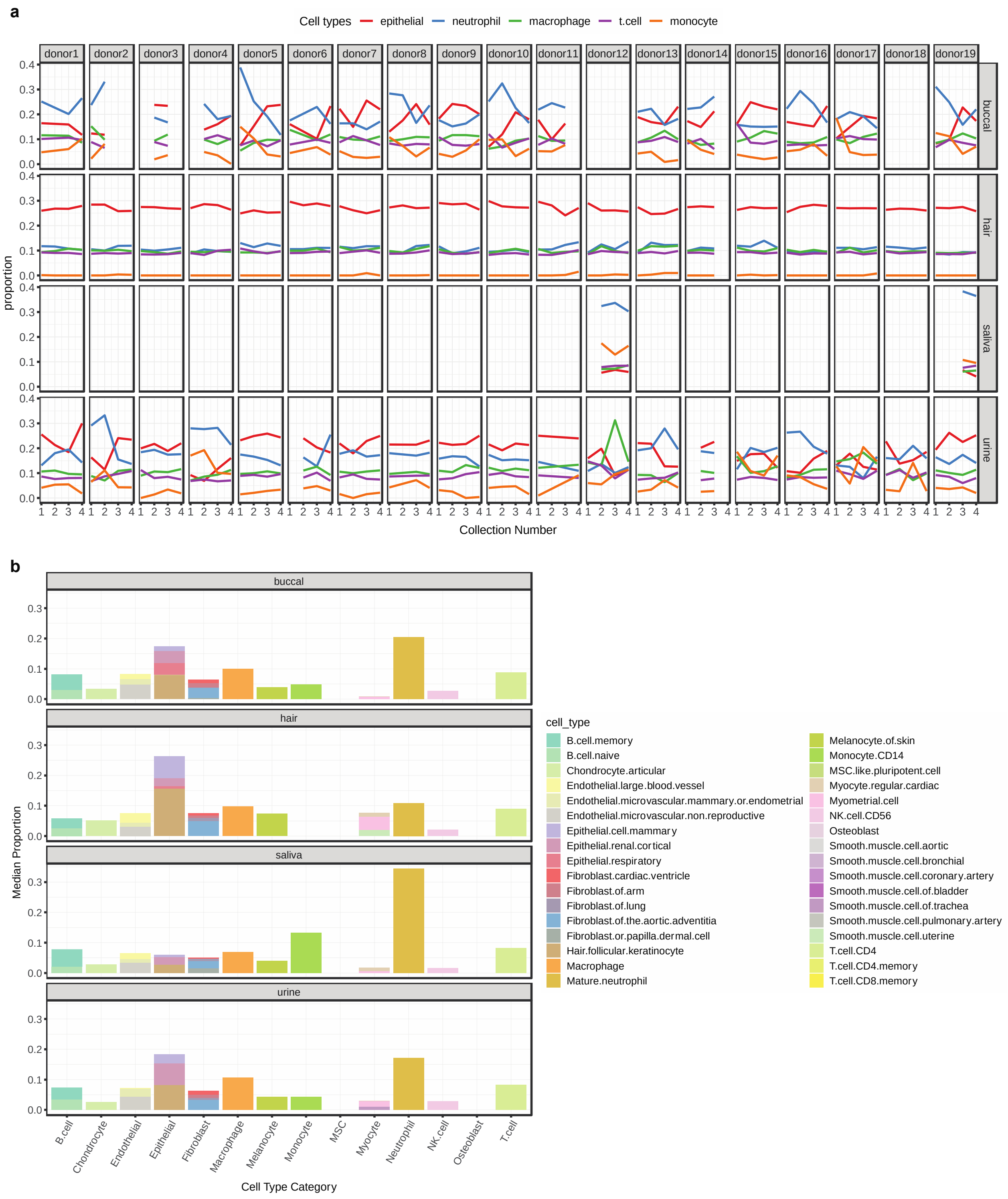
a
Cell types
epithelial
neutrophil
macrophage
t.cell
monocyte
donor1
donor2
donor3
donor4
donor5
donor6
donor7
donor8
donor9
donor10
donor11
donor12
donor13
donor14
donor15
donor16
donor17
donor18
donor19
buccal
hair
saliva
urine
proportion
Collection Number
b
buccal
hair
saliva
urine
Median Proportion
B.cell
Chondrocyte
Endothelial
Epithelial
Fibroblast
Macrophage
Melanocyte
Monocyte
MSC
Myocyte
Neutrophil
NK.cell
Osteoblast
T.cell
Cell Type Category
cell_type
B.cell.memory
B.cell.naive
Chondrocyte.articular
Endothelial.large.blood.vessel
Endothelial.microvascular.mammary.or.endometrial
Endothelial.microvascular.non.reproductive
Epithelial.cell.mammary
Epithelial.renal.cortical
Epithelial.respiratory
Fibroblast.cardiac.ventricle
Fibroblast.of.arm
Fibroblast.of.lung
Fibroblast.of.the.aortic.adventitia
Fibroblast.or.papilla.dermal.cell
Hair.follicular.keratinocyte
Macrophage
Mature.neutrophil
Melanocyte.of.skin
Monocyte.CD14
MSC.like.pluripotent.cell
Myocyte.regular.cardiac
Myometrial.cell
NK.cell.CD56
Osteoblast
Smooth.muscle.cell.aortic
Smooth.muscle.cell.bronchial
Smooth.muscle.cell.coronary.artery
Smooth.muscle.cell.of.bladder
Smooth.muscle.cell.of.trachea
Smooth.muscle.cell.pulmonary.artery
Smooth.muscle.cell.uterine
T.cell.CD4
T.cell.CD4.memory
T.cell.CD8.memory

**Supplementary Figure 7. a.** GEDIT cell type proportion estimates per collection and donor. Top 25% most abundant, condensed cell type categories are shown. **b.** Breakdown of all cell types included in the GEDIT reference. Binning into larger cell type categories is shown.

a

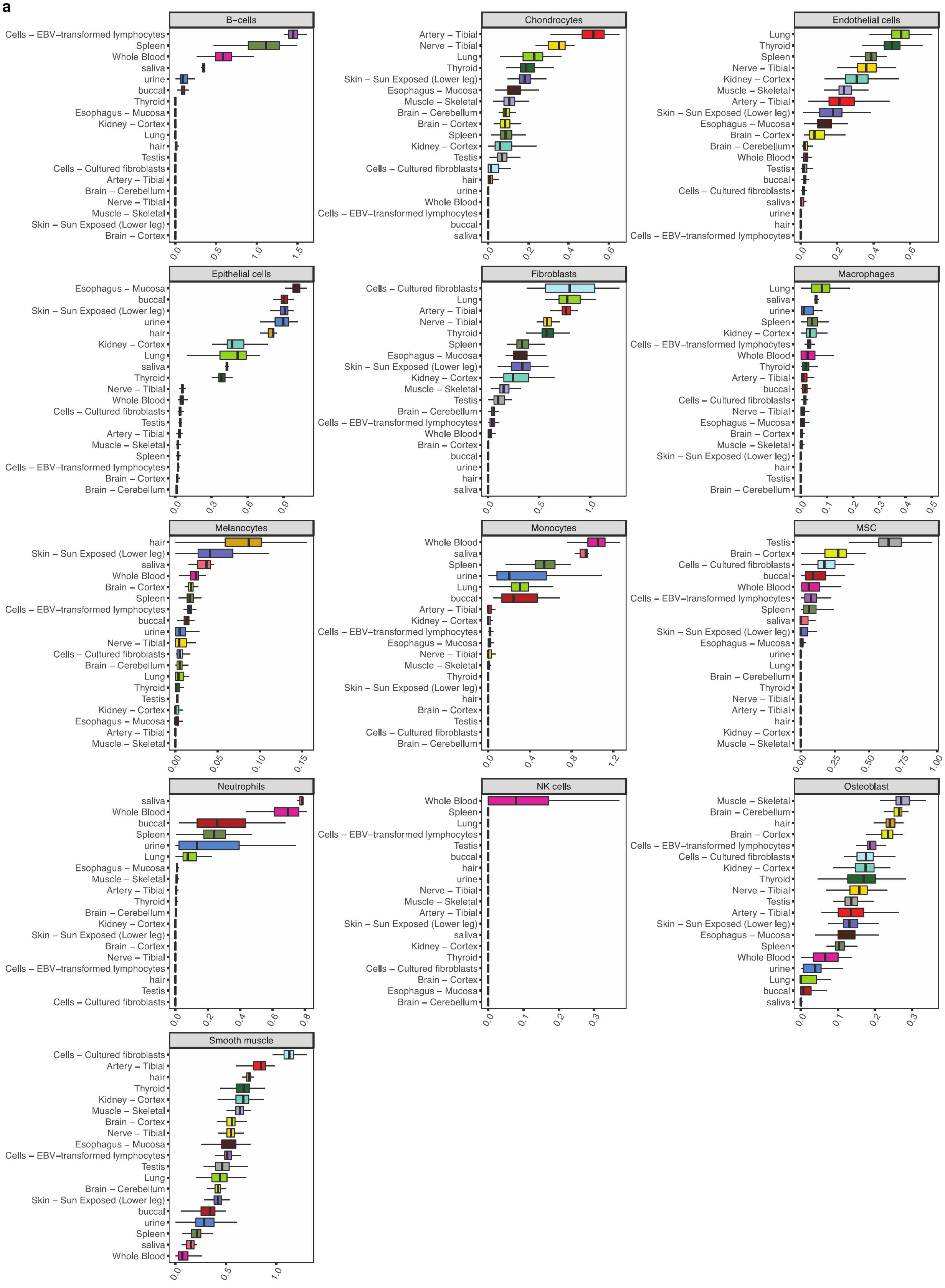
B−cells
Cells − EBV−transformed lymphocytes
Spleen
Whole Blood
saliva
urine
buccal
Thyroid
Esophagus − Mucosa
Kidney − Cortex
Lung
hair
Testis
Cells − Cultured fibroblasts
Artery − Tibial
Brain − Cerebellum
Nerve − Tibial
Muscle − Skeletal
Skin − Sun Exposed (Lower leg)
Brain − Cortex
0.0 0.5 1.0 1.5
Chondrocytes
Artery − Tibial
Nerve − Tibial
Lung
Thyroid
Skin − Sun Exposed (Lower leg)
Esophagus − Mucosa
Muscle − Skeletal
Brain − Cerebellum
Brain − Cortex
Spleen
Kidney − Cortex
Testis
Cells − Cultured fibroblasts
hair
urine
Whole Blood
Cells − EBV−transformed lymphocytes
buccal
saliva
0.0 0.2 0.4 0.6
Endothelial cells
Lung
Thyroid
Spleen
Nerve − Tibial
Kidney − Cortex
Muscle − Skeletal
Artery − Tibial
Skin − Sun Exposed (Lower leg)
Esophagus − Mucosa
Brain − Cortex
Brain − Cerebellum
Whole Blood
Testis
buccal
Cells − Cultured fibroblasts
saliva
urine
hair
Cells − EBV−transformed lymphocytes
0.0 0.2 0.4 0.6
Epithelial cells
Esophagus − Mucosa
buccal
Skin − Sun Exposed (Lower leg)
urine
hair
Kidney − Cortex
Lung
saliva
Thyroid
Nerve − Tibial
Whole Blood
Cells − Cultured fibroblasts
Testis
Artery − Tibial
Muscle − Skeletal
Spleen
Cells − EBV−transformed lymphocytes
Brain − Cortex
Brain − Cerebellum
0.0 0.3 0.6 0.9
Fibroblasts
Cells − Cultured fibroblasts
Lung
Artery − Tibial
Nerve − Tibial
Thyroid
Spleen
Esophagus − Mucosa
Skin − Sun Exposed (Lower leg)
Kidney − Cortex
Muscle − Skeletal
Testis
Brain − Cerebellum
Cells − EBV−transformed lymphocytes
Whole Blood
Brain − Cortex
buccal
urine
hair
saliva
0.0 0.5 1.0
Macrophages
Lung
saliva
urine
Spleen
Kidney − Cortex
Cells − EBV−transformed lymphocytes
Whole Blood
Thyroid
Artery − Tibial
buccal
Cells − Cultured fibroblasts
Nerve − Tibial
Esophagus − Mucosa
Brain − Cortex
Muscle − Skeletal
Skin − Sun Exposed (Lower leg)
hair
Testis
Brain − Cerebellum
0.0 0.1 0.2 0.3 0.4 0.5
Melanocytes
hair
Skin − Sun Exposed (Lower leg)
saliva
Whole Blood
Brain − Cortex
Spleen
Cells − EBV−transformed lymphocytes
buccal
urine
Nerve − Tibial
Cells − Cultured fibroblasts
Brain − Cerebellum
Lung
Thyroid
Testis
Kidney − Cortex
Esophagus − Mucosa
Artery − Tibial
Muscle − Skeletal
0.00 0.05 0.10 0.15
Monocytes
Whole Blood
saliva
Spleen
urine
Lung
buccal
Artery − Tibial
Kidney − Cortex
Cells − EBV−transformed lymphocytes
Esophagus − Mucosa
Nerve − Tibial
Muscle − Skeletal
Thyroid
Skin − Sun Exposed (Lower leg)
hair
Brain − Cortex
Testis
Cells − Cultured fibroblasts
Brain − Cerebellum
0.0 0.4 0.8 1.2
MSC
Testis
Brain − Cortex
Cells − Cultured fibroblasts
buccal
Whole Blood
Cells − EBV−transformed lymphocytes
Spleen
saliva
Skin − Sun Exposed (Lower leg)
Esophagus − Mucosa
urine
Lung
Brain − Cerebellum
Thyroid
Nerve − Tibial
Artery − Tibial
hair
Kidney − Cortex
Muscle − Skeletal
0.00 0.25 0.50 0.75 1.00
Neutrophils
saliva
Whole Blood
buccal
Spleen
urine
Lung
Esophagus − Mucosa
Muscle − Skeletal
Artery − Tibial
Thyroid
Brain − Cerebellum
Kidney − Cortex
Skin − Sun Exposed (Lower leg)
Brain − Cortex
Nerve − Tibial
Cells − EBV−transformed lymphocytes
hair
Testis
Cells − Cultured fibroblasts
0.0 0.2 0.4 0.6 0.8
NK cells
Whole Blood
Spleen
Lung
Cells − EBV−transformed lymphocytes
Testis
buccal
hair
urine
Nerve − Tibial
Muscle − Skeletal
Artery − Tibial
Skin − Sun Exposed (Lower leg)
saliva
Kidney − Cortex
Thyroid
Cells − Cultured fibroblasts
Brain − Cortex
Esophagus − Mucosa
Brain − Cerebellum
0.0 0.1 0.2 0.3
Osteoblast
Muscle − Skeletal
Brain − Cerebellum
hair
Brain − Cortex
Cells − EBV−transformed lymphocytes
Cells − Cultured fibroblasts
Kidney − Cortex
Thyroid
Nerve − Tibial
Testis
Artery − Tibial
Skin − Sun Exposed (Lower leg)
Esophagus − Mucosa
Spleen
Whole Blood
urine
Lung
buccal
saliva
0.0 0.1 0.2 0.3
Smooth muscle
Cells − Cultured fibroblasts
Artery − Tibial
hair
Thyroid
Kidney − Cortex
Muscle − Skeletal
Brain − Cortex
Nerve − Tibial
Esophagus − Mucosa
Cells − EBV−transformed lymphocytes
Testis
Lung
Brain − Cerebellum
Skin − Sun Exposed (Lower leg)
buccal
urine
Spleen
saliva
Whole Blood
0.0 0.5 1.0

**b**

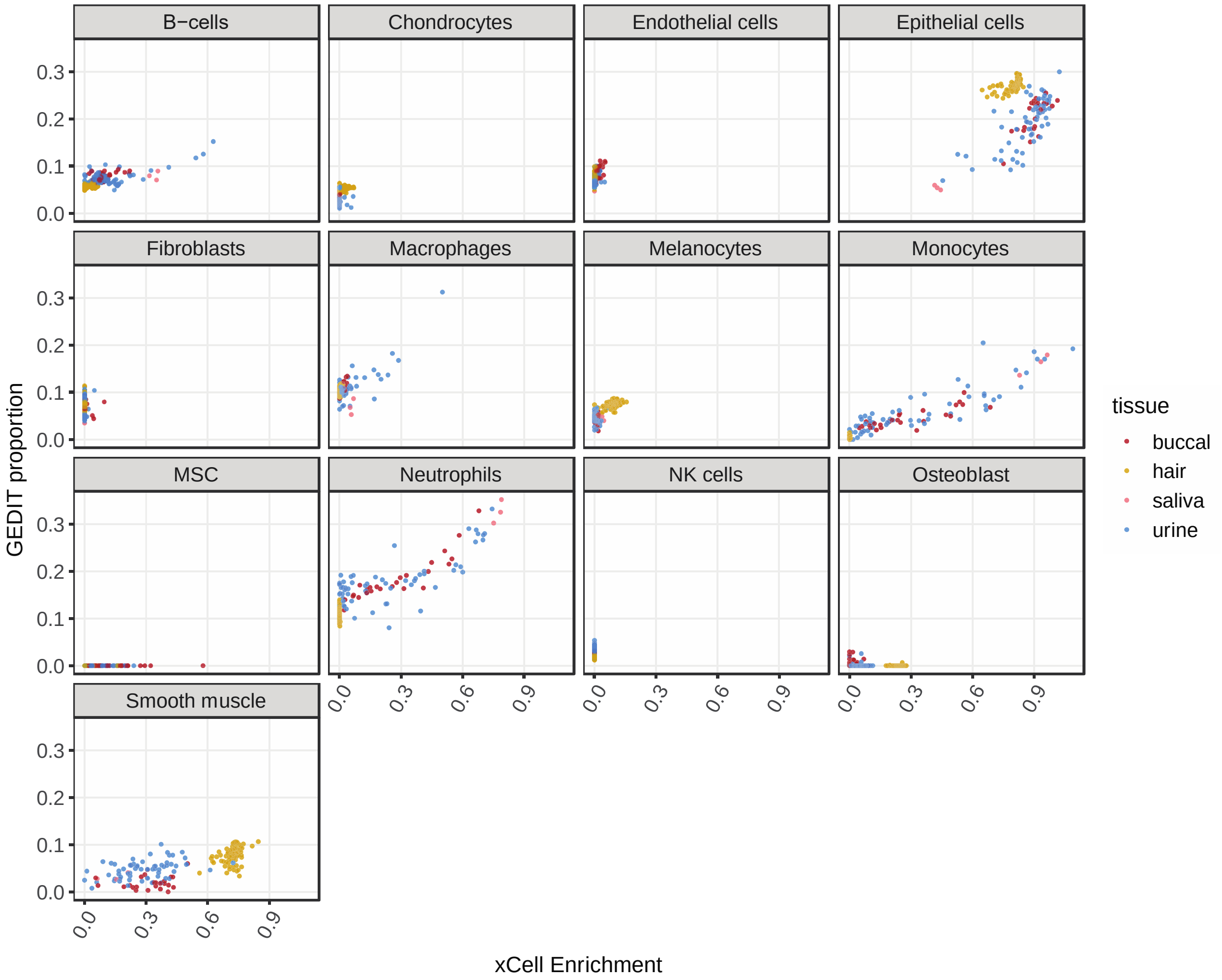


**Supplementary Figure 8. a.** xCell enrichment scores across the noninvasive and select GTEx tissues for cell types corresponding to the GEDIT collapsed cell type categories. Note that the enrichment score does not correspond to a proportion. **b.** Comparison of xCell enrichment scores and GEDIT proportions for cell types shared by both references. Only noninvasive tissues are shown.

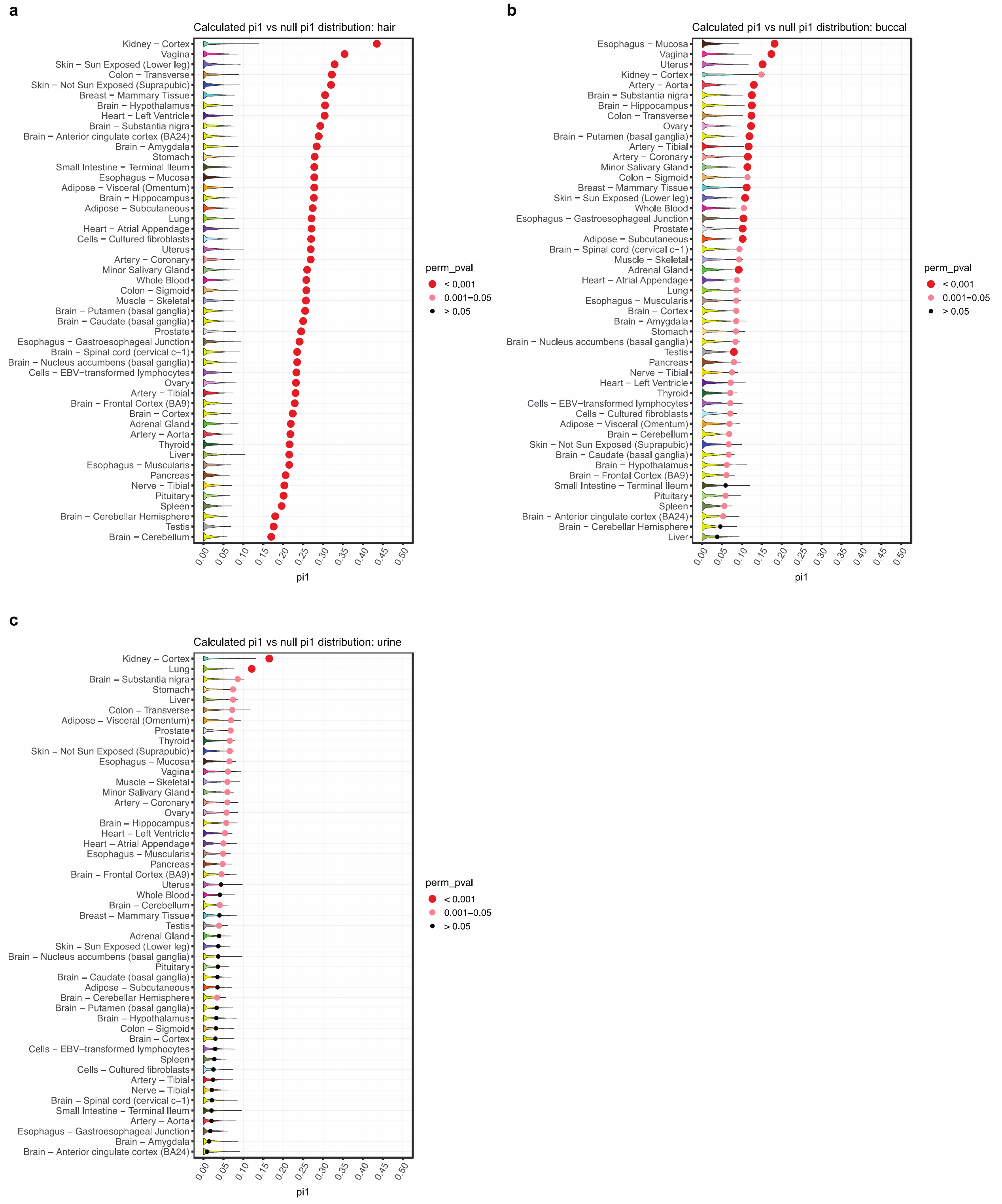
a
Calculated pi1 vs null pi1 distribution: hair
Kidney – Cortex
Vagina
Skin – Sun Exposed (Lower leg)
Colon – Transverse
Skin – Not Sun Exposed (Suprapubic)
Breast – Mammary Tissue
Brain – Hypothalamus
Heart – Left Ventricle
Brain – Substantia nigra
Brain – Anterior cingulate cortex (BA24)
Brain – Amygdala
Stomach
Small Intestine – Terminal Ileum
Esophagus – Mucosa
Adipose – Visceral (Omentum)
Brain – Hippocampus
Adipose – Subcutaneous
Lung
Heart – Atrial Appendage
Cells – Cultured fibroblasts
Uterus
Artery – Coronary
Minor Salivary Gland
Whole Blood
Colon – Sigmoid
Muscle – Skeletal
Brain – Putamen (basal ganglia)
Brain – Caudate (basal ganglia)
Prostate
Esophagus – Gastroesophageal Junction
Brain – Spinal cord (cervical c-1)
Brain – Nucleus accumbens (basal ganglia)
Cells – EBV-transformed lymphocytes
Ovary
Artery – Tibial
Brain – Frontal Cortex (BA9)
Brain – Cortex
Adrenal Gland
Artery – Aorta
Thyroid
Liver
Esophagus – Muscularis
Pancreas
Nerve – Tibial
Pituitary
Spleen
Brain – Cerebellar Hemisphere
Testis
Brain – Cerebellum
0.00 0.05 0.10 0.15 0.20 0.25 0.30 0.35 0.40 0.45 0.50
pi1
perm_pval
< 0.001
0.001–0.05
> 0.05
b
Calculated pi1 vs null pi1 distribution: buccal
Esophagus – Mucosa
Vagina
Uterus
Kidney – Cortex
Artery – Aorta
Brain – Substantia nigra
Brain – Hippocampus
Colon – Transverse
Ovary
Brain – Putamen (basal ganglia)
Artery – Tibial
Artery – Coronary
Minor Salivary Gland
Colon – Sigmoid
Breast – Mammary Tissue
Skin – Sun Exposed (Lower leg)
Whole Blood
Esophagus – Gastroesophageal Junction
Prostate
Adipose – Subcutaneous
Brain – Spinal cord (cervical c-1)
Muscle – Skeletal
Adrenal Gland
Heart – Atrial Appendage
Lung
Esophagus – Muscularis
Brain – Cortex
Brain – Amygdala
Stomach
Brain – Nucleus accumbens (basal ganglia)
Testis
Pancreas
Nerve – Tibial
Heart – Left Ventricle
Thyroid
Cells – EBV-transformed lymphocytes
Cells – Cultured fibroblasts
Adipose – Visceral (Omentum)
Brain – Cerebellum
Skin – Not Sun Exposed (Suprapubic)
Brain – Caudate (basal ganglia)
Brain – Hypothalamus
Brain – Frontal Cortex (BA9)
Small Intestine – Terminal Ileum
Pituitary
Spleen
Brain – Anterior cingulate cortex (BA24)
Brain – Cerebellar Hemisphere
Liver
0.00 0.05 0.10 0.15 0.20 0.25 0.30 0.35 0.40 0.45 0.50
pi1
perm_pval
< 0.001
0.001–0.05
> 0.05
c
Calculated pi1 vs null pi1 distribution: urine
Kidney – Cortex
Lung
Brain – Substantia nigra
Stomach
Liver
Colon – Transverse
Adipose – Visceral (Omentum)
Prostate
Thyroid
Skin – Not Sun Exposed (Suprapubic)
Esophagus – Mucosa
Vagina
Muscle – Skeletal
Minor Salivary Gland
Artery – Coronary
Ovary
Brain – Hippocampus
Heart – Left Ventricle
Heart – Atrial Appendage
Esophagus – Muscularis
Pancreas
Brain – Frontal Cortex (BA9)
Uterus
Whole Blood
Brain – Cerebellum
Breast – Mammary Tissue
Testis
Adrenal Gland
Skin – Sun Exposed (Lower leg)
Brain – Nucleus accumbens (basal ganglia)
Pituitary
Brain – Caudate (basal ganglia)
Adipose – Subcutaneous
Brain – Cerebellar Hemisphere
Brain – Putamen (basal ganglia)
Brain – Hypothalamus
Colon – Sigmoid
Brain – Cortex
Cells – EBV-transformed lymphocytes
Spleen
Cells – Cultured fibroblasts
Artery – Tibial
Nerve – Tibial
Brain – Spinal cord (cervical c-1)
Small Intestine – Terminal Ileum
Artery – Aorta
Esophagus – Gastroesophageal Junction
Brain – Amygdala
Brain – Anterior cingulate cortex (BA24)
0.00 0.05 0.10 0.15 0.20 0.25 0.30 0.35 0.40 0.45 0.50
pi1
perm_pval
< 0.001
0.001–0.05
> 0.05

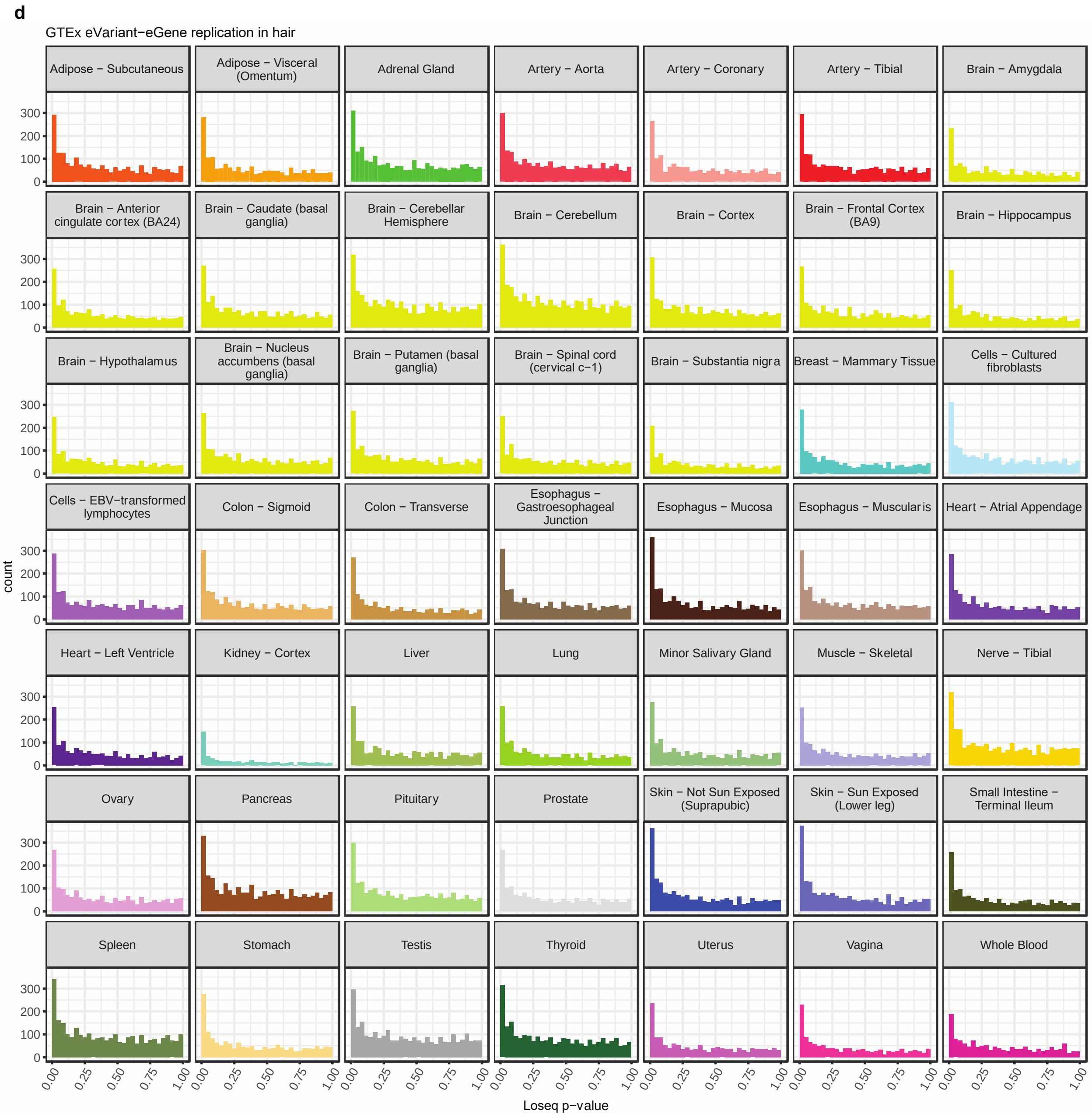
d
GTEx eVariant−eGene replication in hair
Adipose − Subcutaneous
Adipose − Visceral (Omentum)
Adrenal Gland
Artery − Aorta
Artery − Coronary
Artery − Tibial
Brain − Amygdala
Brain − Anterior cingulate cortex (BA24)
Brain − Caudate (basal ganglia)
Brain − Cerebellar Hemisphere
Brain − Cerebellum
Brain − Cortex
Brain − Frontal Cortex (BA9)
Brain − Hippocampus
Brain − Hypothalamus
Brain − Nucleus accumbens (basal ganglia)
Brain − Putamen (basal ganglia)
Brain − Spinal cord (cervical c−1)
Brain − Substantia nigra
Breast − Mammary Tissue
Cells − Cultured fibroblasts
Cells − EBV−transformed lymphocytes
Colon − Sigmoid
Colon − Transverse
Esophagus − Gastroesophageal Junction
Esophagus − Mucosa
Esophagus − Muscularis
Heart − Atrial Appendage
Heart − Left Ventricle
Kidney − Cortex
Liver
Lung
Minor Salivary Gland
Muscle − Skeletal
Nerve − Tibial
Ovary
Pancreas
Pituitary
Prostate
Skin − Not Sun Exposed (Suprapubic)
Skin − Sun Exposed (Lower leg)
Small Intestine − Terminal Ileum
Spleen
Stomach
Testis
Thyroid
Uterus
Vagina
Whole Blood
count
0
100
200
300
0.00
0.25
0.50
0.75
1.00
Loseq p−value

**e**

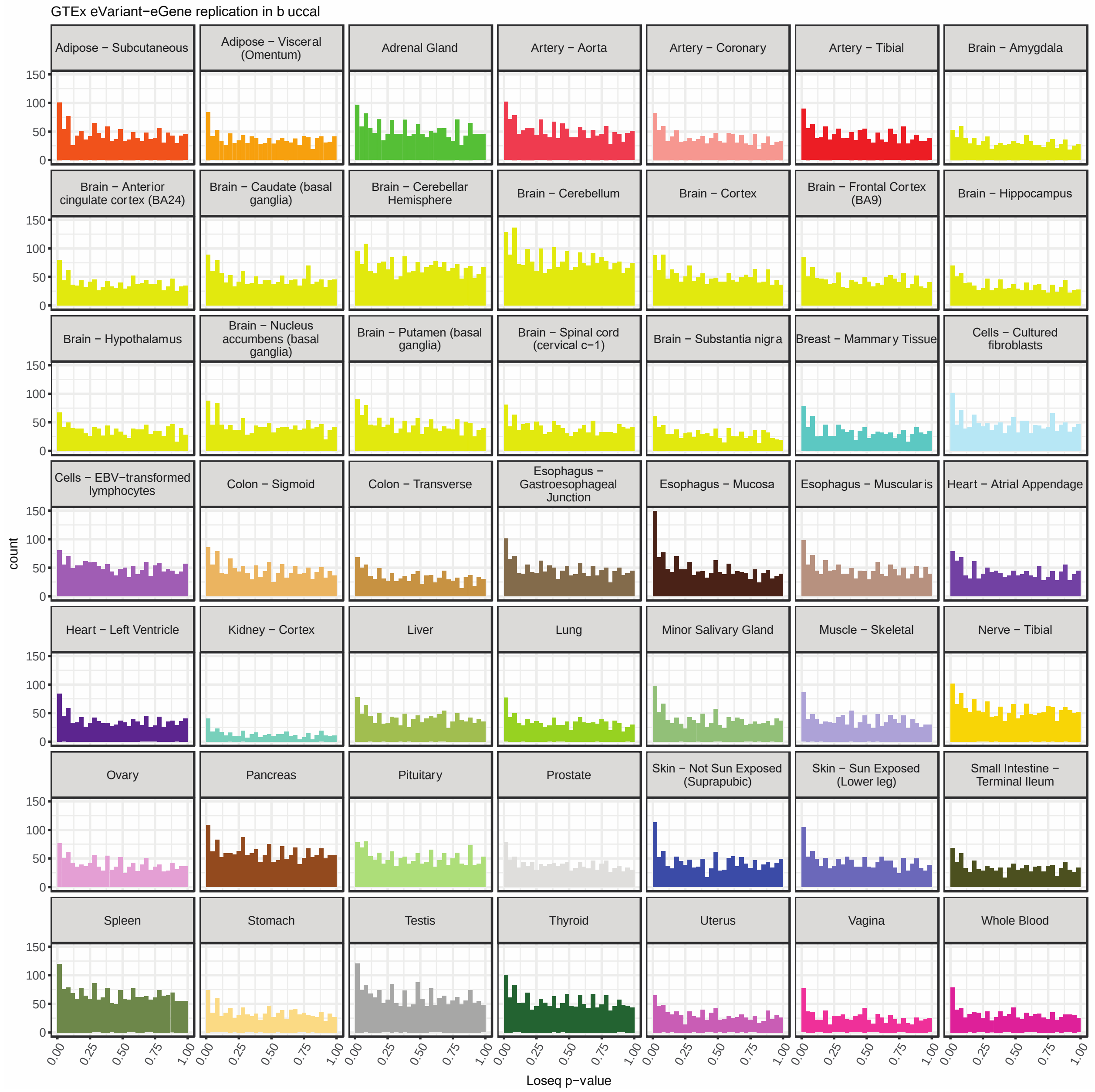

GTEx eVariant−eGene replication in b uccal
Adipose − Subcutaneous
Adipose − Visceral (Omentum)
Adrenal Gland
Artery − Aorta
Artery − Coronary
Artery − Tibial
Brain − Amygdala
Brain − Anterior cingulate cortex (BA24)
Brain − Caudate (basal ganglia)
Brain − Cerebellar Hemisphere
Brain − Cerebellum
Brain − Cortex
Brain − Frontal Cortex (BA9)
Brain − Hippocampus
Brain − Hypothalamus
Brain − Nucleus accumbens (basal ganglia)
Brain − Putamen (basal ganglia)
Brain − Spinal cord (cervical c−1)
Brain − Substantia nigra
Breast − Mammary Tissue
Cells − Cultured fibroblasts
Cells − EBV−transformed lymphocytes
Colon − Sigmoid
Colon − Transverse
Esophagus − Gastroesophageal Junction
Esophagus − Mucosa
Esophagus − Muscularis
Heart − Atrial Appendage
Heart − Left Ventricle
Kidney − Cortex
Liver
Lung
Minor Salivary Gland
Muscle − Skeletal
Nerve − Tibial
Ovary
Pancreas
Pituitary
Prostate
Skin − Not Sun Exposed (Suprapubic)
Skin − Sun Exposed (Lower leg)
Small Intestine − Terminal Ileum
Spleen
Stomach
Testis
Thyroid
Uterus
Vagina
Whole Blood
count
0
50
100
150
0.00
0.25
0.50
0.75
1.00
Loseq p−value

**f**

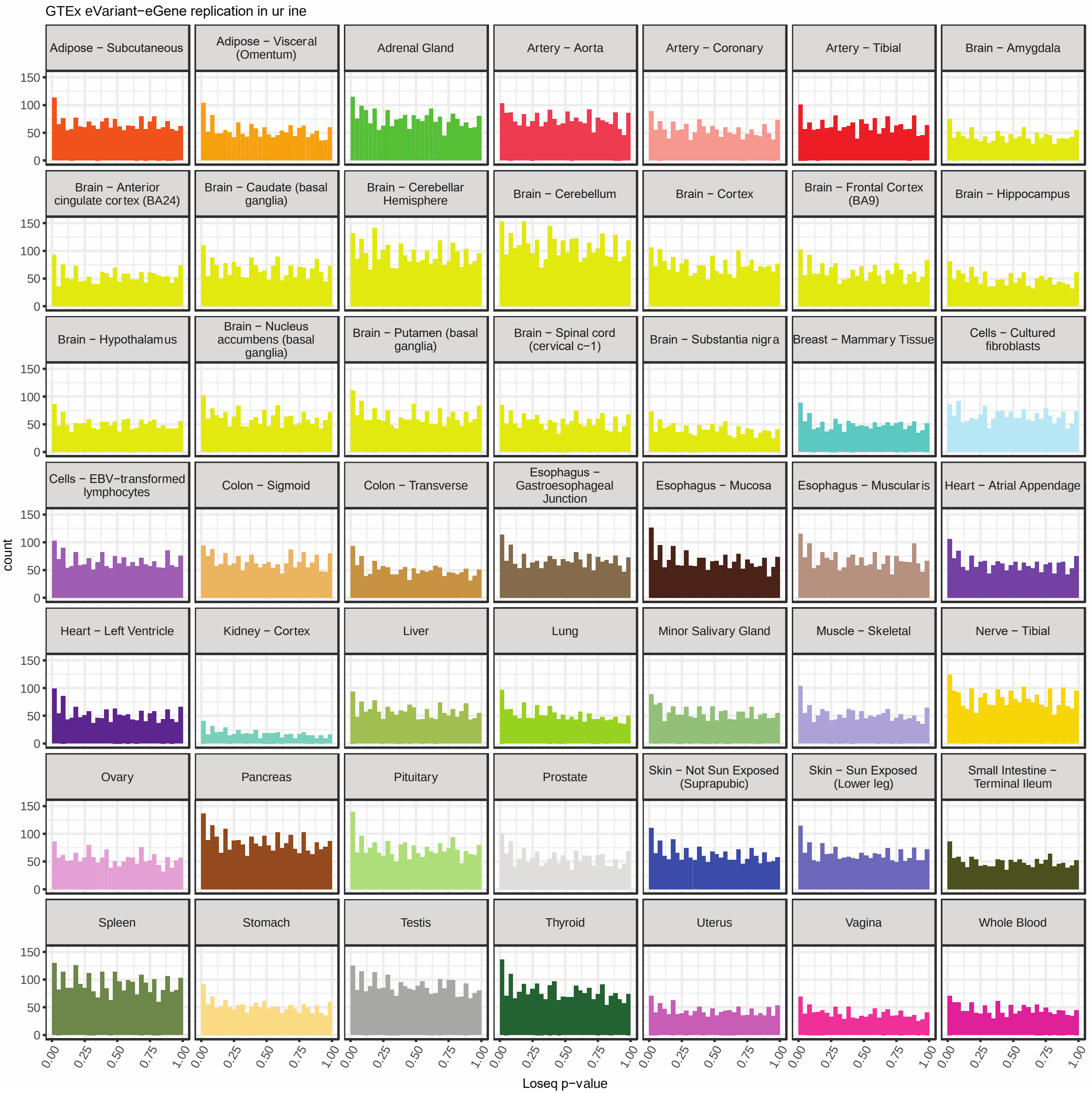


**Supplementary Figure 9. a. b. c.** Calculated pi1 versus the null pi1 distribution for every tissue in GTEx. Pi1 was calculated by selecting for significant variants (q-value <=0.05) with MAF > 0.05 and minimum effect size greater than the maximum minimum across GTEx tissues (kidney cortex 0.32) that were present in the noninvasive dataset. The null distribution was generated by performing 1000 samples of size equivalent to the number of overlapping gene-variant pairs used for the pi1 calculation for that tissue. **d. e. f.** Histograms of Loseq p-values for gene-variant pairs included in the pi1 calculation.

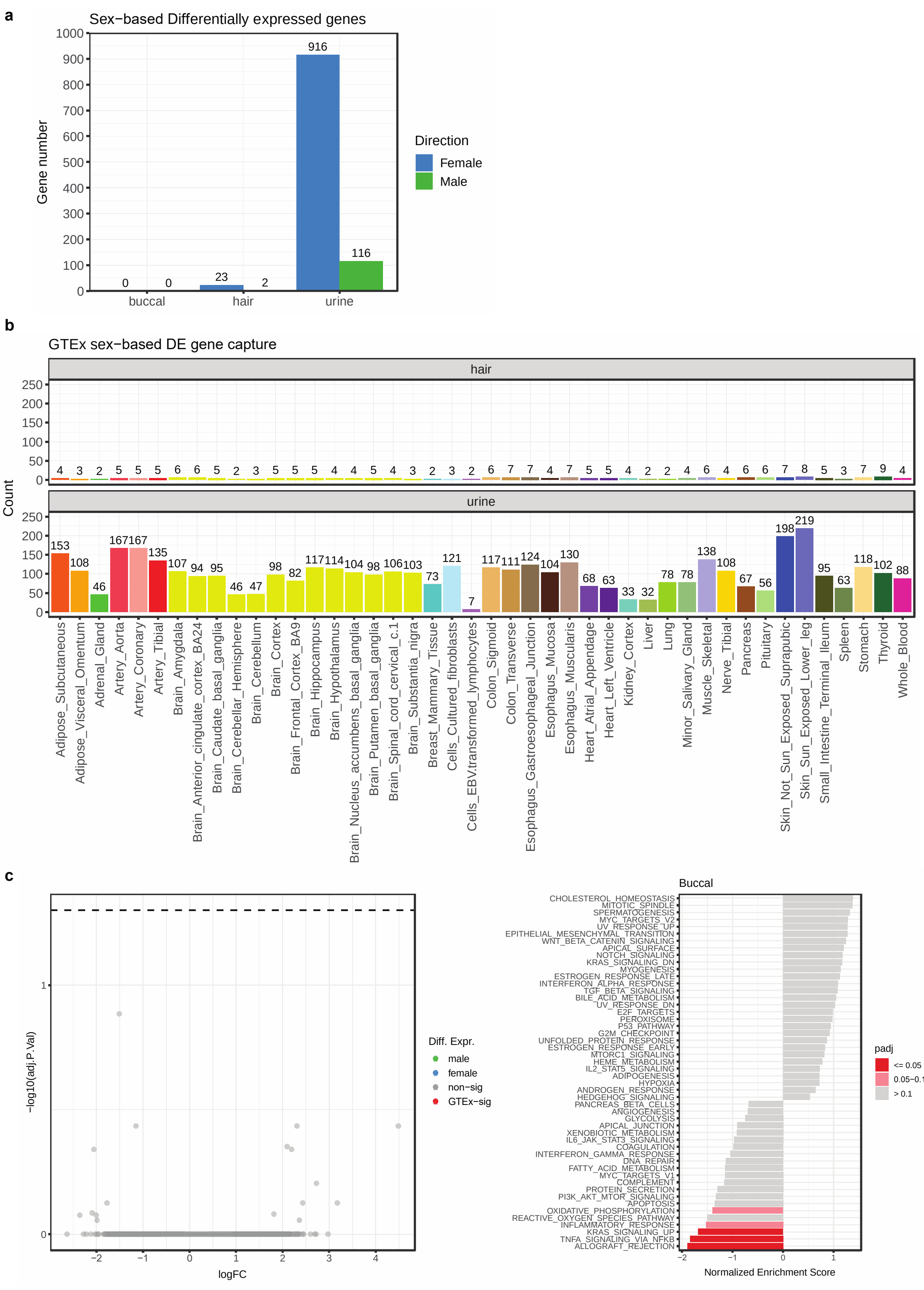
a
Sex−based Differentially expressed genes
Gene number
1000
900
800
700
600
500
400
300
200
100
0
0 0
23 2
916 116
buccal
hair
urine
Direction
Female
Male
b
GTEx sex−based DE gene capture
hair
urine
Count
250
200
150
100
50
0
4 3 2 5 5 5 6 6 5 2 3 5 5 5 4 4 5 4 3 2 3 2 6 7 7 4 7 5 5 4 2 2 4 6 4 6 6 7 8 5 3 7 9 4
153 108 46 167 167 135 107 94 95 46 47 98 82 117 114 104 98 106 103 73 121 7 117 111 124 104 130 68 63 33 32 78 78 138 108 67 56 198 219 95 63 118 102 88
Adipose_Subcutaneous
Adipose_Visceral_Omentum
Adrenal_Gland
Artery_Aorta
Artery_Coronary
Artery_Tibial
Brain_Amygdala
Brain_Anterior_cingulate_cortex_BA24
Brain_Caudate_basal_ganglia
Brain_Cerebellar_Hemisphere
Brain_Cerebellum
Brain_Cortex
Brain_Frontal_Cortex_BA9
Brain_Hippocampus
Brain_Hypothalamus
Brain_Nucleus_accumbens_basal_ganglia
Brain_Putamen_basal_ganglia
Brain_Spinal_cord_cervical_c.1
Brain_Substantia_nigra
Breast_Mammary_Tissue
Cells_Cultured_fibroblasts
Cells_EBV.transformed_lymphocytes
Colon_Sigmoid
Colon_Transverse
Esophagus_Gastroesophageal_Junction
Esophagus_Mucosa
Esophagus_Muscularis
Heart_Atrial_Appendage
Heart_Left_Ventricle
Kidney_Cortex
Liver
Lung
Minor_Salivary_Gland
Muscle_Skeletal
Nerve_Tibial
Pancreas
Pituitary
Skin_Not_Sun_Exposed_Suprapubic
Skin_Sun_Exposed_Lower_leg
Small_Intestine_Terminal_Ileum
Spleen
Stomach
Thyroid
Whole_Blood
c
−log10(adj.P.Val)
1
0
logFC
−2 −1 0 1 2 3 4
Diff. Expr.
male
female
non−sig
GTEx−sig
Buccal
CHOLESTEROL_HOMEOSTASIS
MITOTIC_SPINDLE
SPERMATOGENESIS
MYC_TARGETS_V2
UV_RESPONSE_UP
EPITHELIAL_MESENCHYMAL_TRANSITION
WNT_BETA_CATENIN_SIGNALING
APICAL_SURFACE
NOTCH_SIGNALING
KRAS_SIGNALING_DN
MYOGENESIS
ESTROGEN_RESPONSE_LATE
INTERFERON_ALPHA_RESPONSE
TGF_BETA_SIGNALING
BILE_ACID_METABOLISM
UV_RESPONSE_DN
E2F_TARGETS
PEROXISOME
P53_PATHWAY
G2M_CHECKPOINT
UNFOLDED_PROTEIN_RESPONSE
ESTROGEN_RESPONSE_EARLY
MTORC1_SIGNALING
HEME_METABOLISM
IL2_STAT5_SIGNALING
ADIPOGENESIS
HYPOXIA
ANDROGEN_RESPONSE
HEDGEHOG_SIGNALING
PANCREAS_BETA_CELLS
ANGIOGENESIS
GLYCOLYSIS
APICAL_JUNCTION
XENOBIOTIC_METABOLISM
IL6_JAK_STAT3_SIGNALING
COAGULATION
INTERFERON_GAMMA_RESPONSE
DNA_REPAIR
FATTY_ACID_METABOLISM
MYC_TARGETS_V1
COMPLEMENT
PROTEIN_SECRETION
PI3K_AKT_MTOR_SIGNALING
APOPTOSIS
OXIDATIVE_PHOSPHORYLATION
REACTIVE_OXYGEN_SPECIES_PATHWAY
INFLAMMATORY_RESPONSE
KRAS_SIGNALING_UP
TNFA_SIGNALING_VIA_NFKB
ALLOGRAFT_REJECTION
−2 −1 0 1
Normalized Enrichment Score
padj
<= 0.05
0.05−0.1
> 0.1

**Supplementary Figure 10. a.** Number of sex-based upregulated genes per noninvasive tissue type. **b.** Per tissue overlap between significant DE genes in the noninvasive dataset with genes previously found to be significant in the GTEx dataset. **c.** Sex-based differential expression for buccal samples, with no significant genes. FGSEA shows some rank-based gene category enrichment for females.

**a**

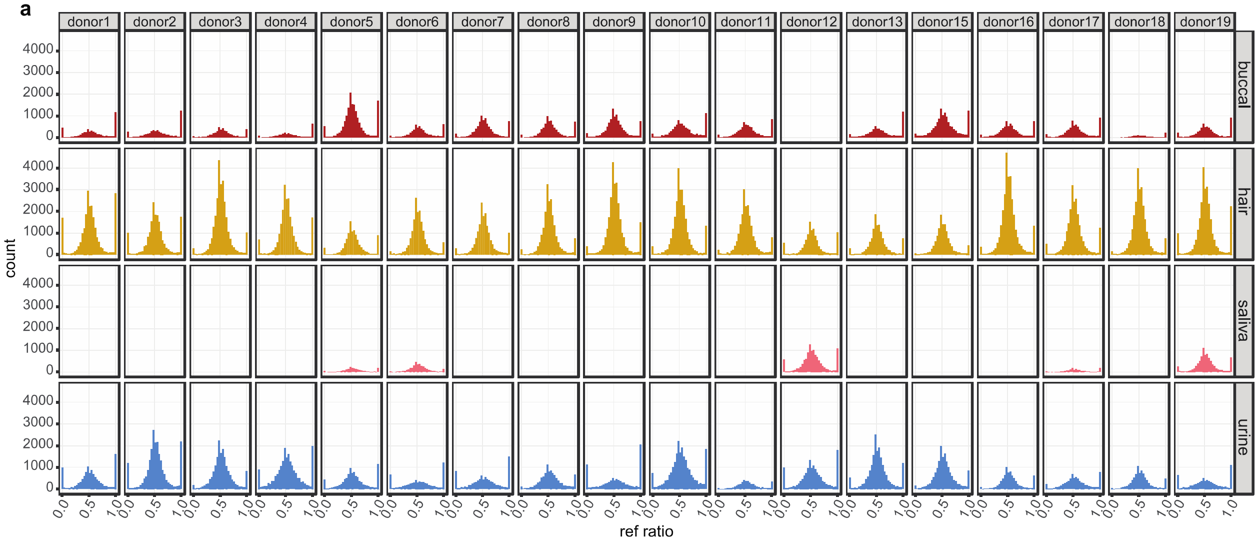

donor1
donor2
donor3
donor4
donor5
donor6
donor7
donor8
donor9
donor10
donor11
donor12
donor13
donor15
donor16
donor17
donor18
donor19
buccal
hair
saliva
urine
count
0
1000
2000
3000
4000
0.0
0.5
1.0
ref ratio


**b**

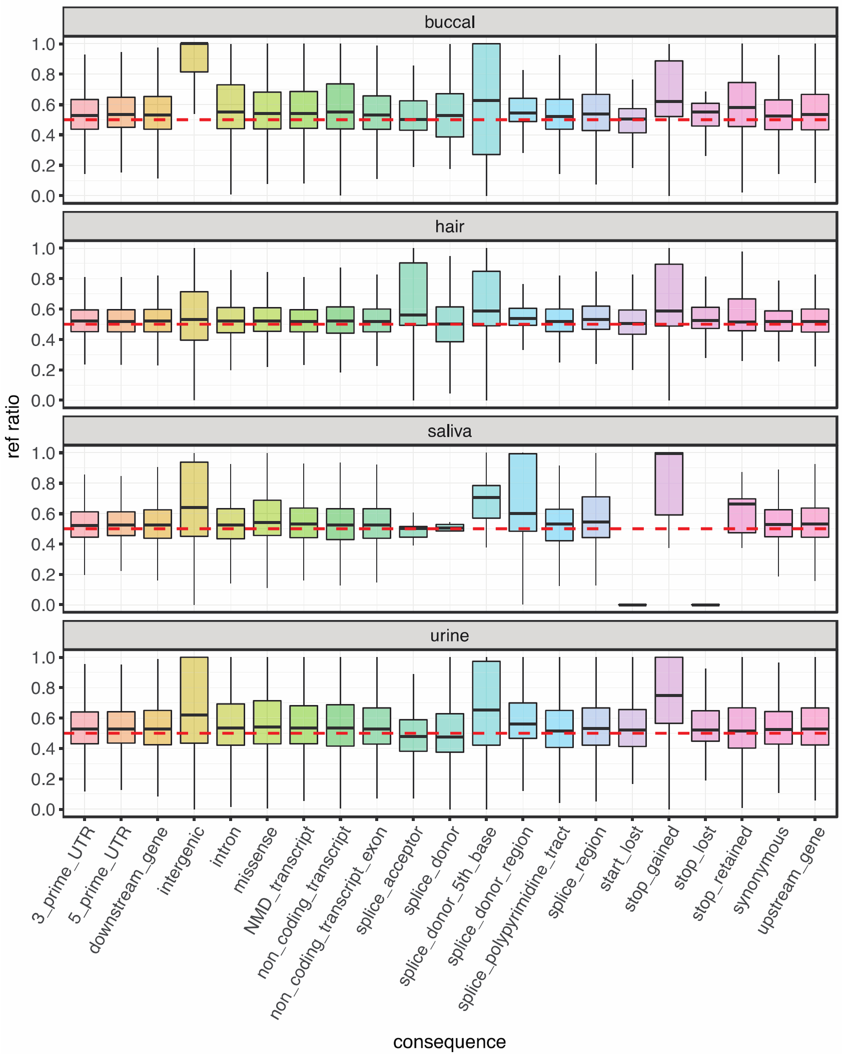

buccal
hair
saliva
urine
ref ratio
0.0
0.2
0.4
0.6
0.8
1.0
3_prime_UTR
5_prime_UTR
downstream_gene
intergenic
intron
missense
NMD_transcript
non_coding_transcript
non_coding_transcript_exon
splice_acceptor
splice_donor
splice_donor_5th_base
splice_donor_region
splice_polypyrimidine_tract
splice_region
start_lost
stop_gained
stop_lost
stop_retained
synonymous
upstream_gene
consequence

**Supplementary Figure 11. a.** Ratio of (reference allele count)/(total count) for all heterozygous sites per donor and tissue. **b.** Reference ratio breakdown per VEP annotation.

a

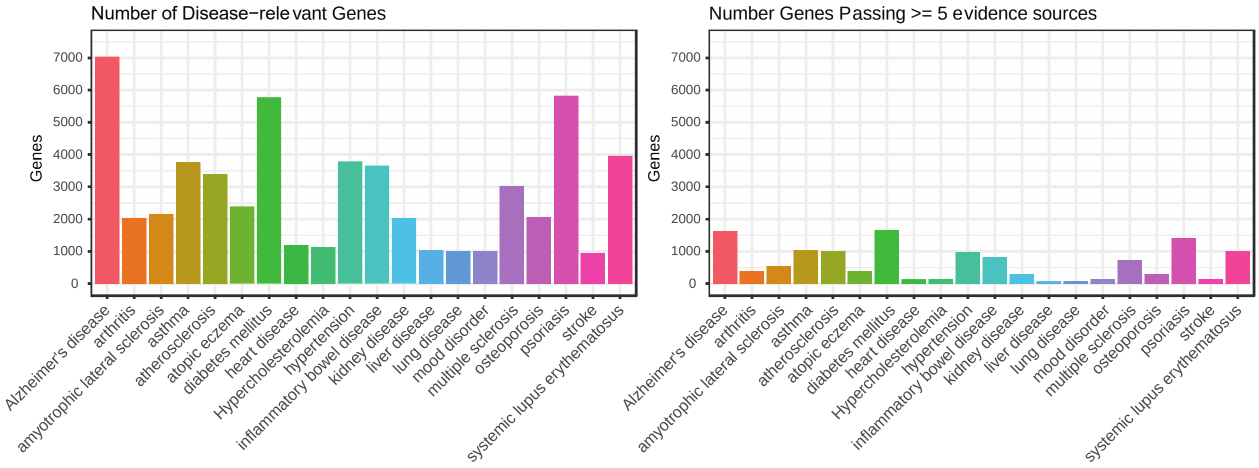
Number of Disease−relevant Genes
Genes
Number Genes Passing >= 5 evidence sources
Genes
Alzheimer's disease
arthritis
amyotrophic lateral sclerosis
asthma
atherosclerosis
atopic eczema
diabetes mellitus
heart disease
Hypercholesterolemia
hypertension
inflammatory bowel disease
kidney disease
liver disease
lung disease
mood disorder
multiple sclerosis
osteoporosis
psoriasis
stroke
systemic lupus erythematosus

b

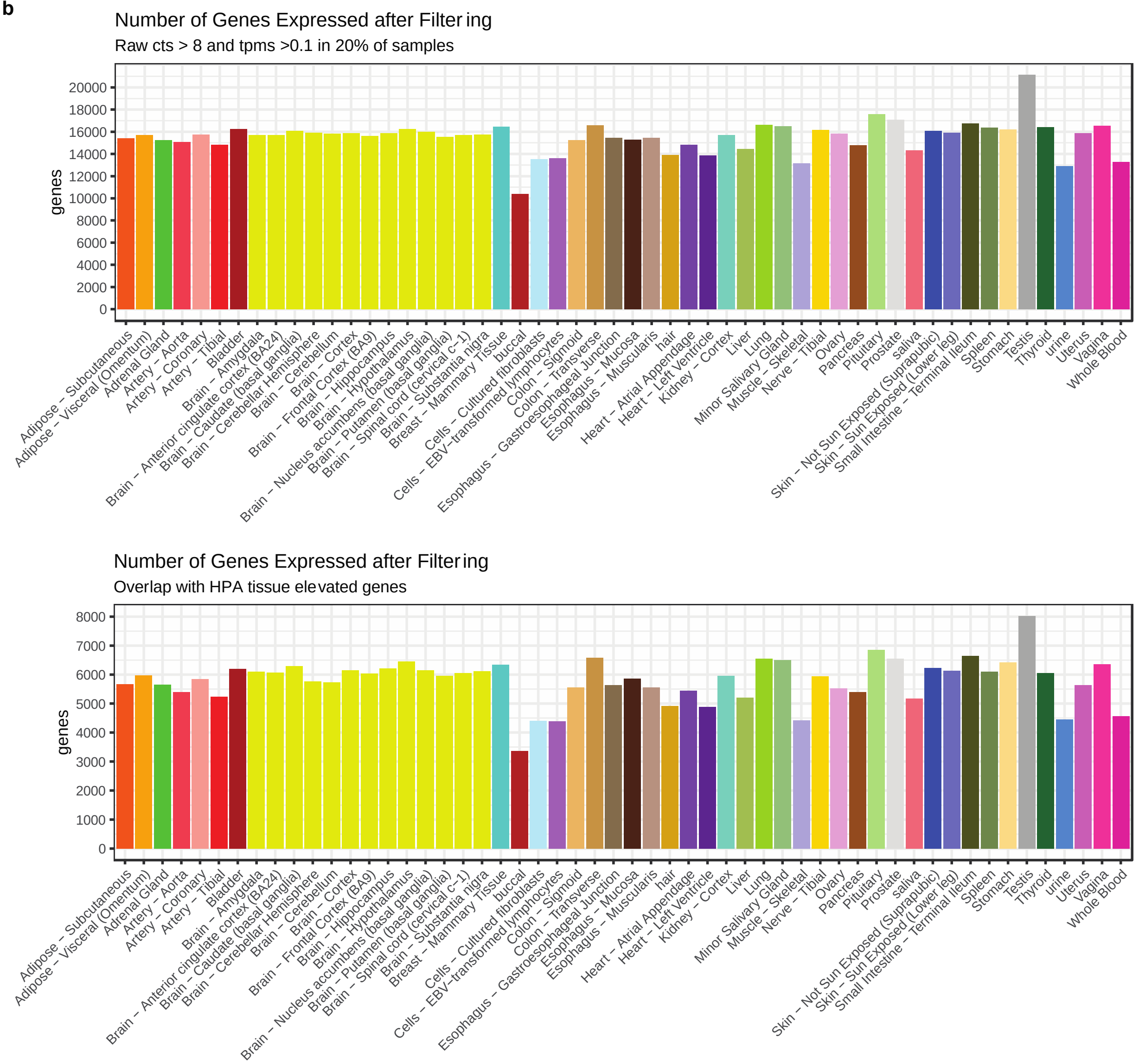
Number of Genes Expressed after Filtering
Raw cts > 8 and tpms >0.1 in 20% of samples
genes
Number of Genes Expressed after Filtering
Overlap with HPA tissue elevated genes
genes
Adipose − Subcutaneous
Adipose − Visceral (Omentum)
Adrenal Gland
Artery − Aorta
Artery − Coronary
Artery − Tibial
Bladder
Brain − Amygdala
Brain − Anterior cingulate cortex (BA24)
Brain − Caudate (basal ganglia)
Brain − Cerebellar Hemisphere
Brain − Cerebellum
Brain − Cortex
Brain − Frontal Cortex (BA9)
Brain − Hippocampus
Brain − Hypothalamus
Brain − Nucleus accumbens (basal ganglia)
Brain − Putamen (basal ganglia)
Brain − Spinal cord (cervical c−1)
Brain − Substantia nigra
Breast − Mammary Tissue
buccal
Cells − Cultured fibroblasts
Cells − EBV−transformed lymphocytes
Colon − Sigmoid
Colon − Transverse
Esophagus − Gastroesophageal Junction
Esophagus − Mucosa
Esophagus − Muscularis
hair
Heart − Atrial Appendage
Heart − Left Ventricle
Kidney − Cortex
Liver
Lung
Minor Salivary Gland
Muscle − Skeletal
Nerve − Tibial
Ovary
Pancreas
Pituitary
Prostate
saliva
Skin − Not Sun Exposed (Suprapubic)
Skin − Sun Exposed (Lower leg)
Small Intestine − Terminal Ileum
Spleen
Stomach
Testis
Thyroid
urine
Uterus
Vagina
Whole Blood

**c**

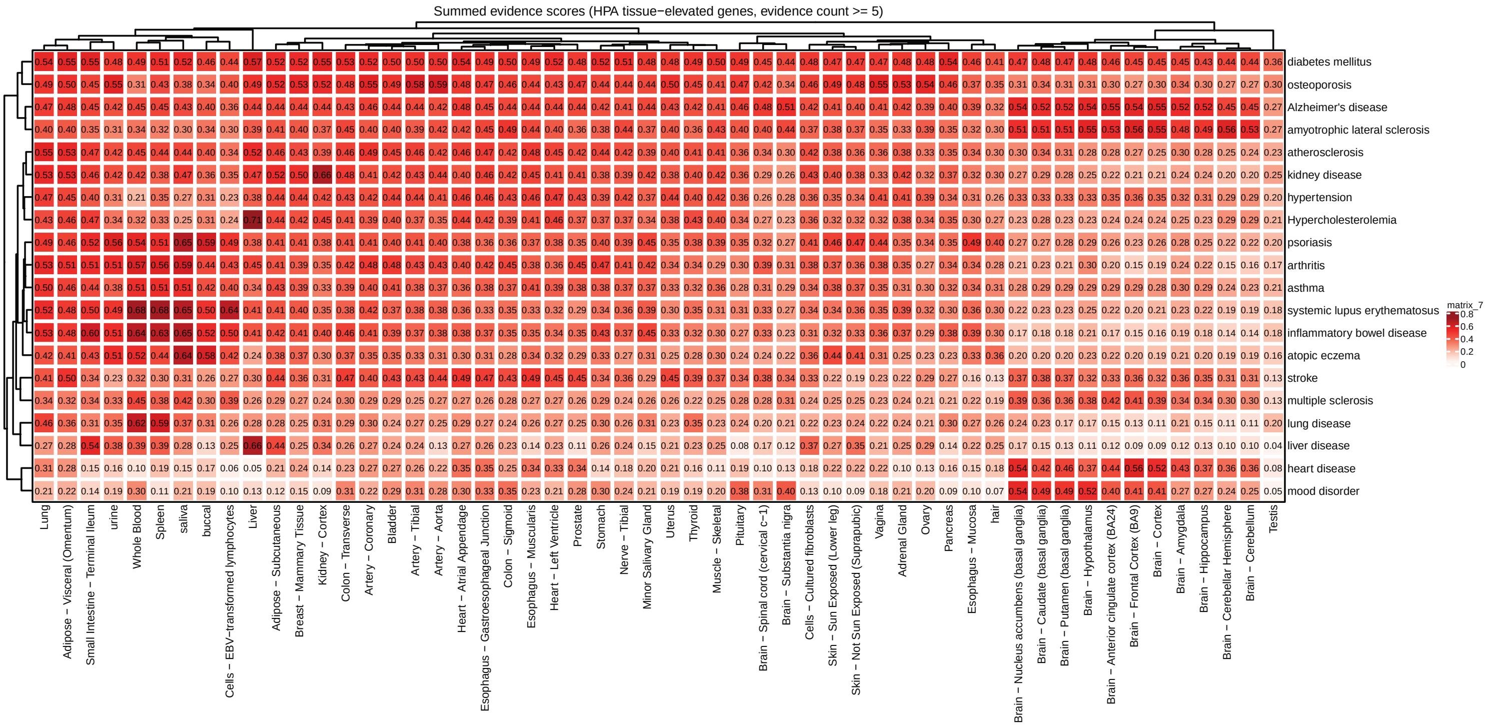


**Supplementary Figure 12. a.** Total number of disease-relevant genes per OpenTargets ontology category. Genes with >= 5 separate sources of evidence were included in the final analysis. **b.** Number of genes per tissue following minimum expression level thresholding and overlap with the HPA tissue-elevated gene list. The top 3,411 most expressed genes per tissue were included in the analysis. **c.** Summed evidence scores (SESs) for all GTEx and noninvasive tissues.

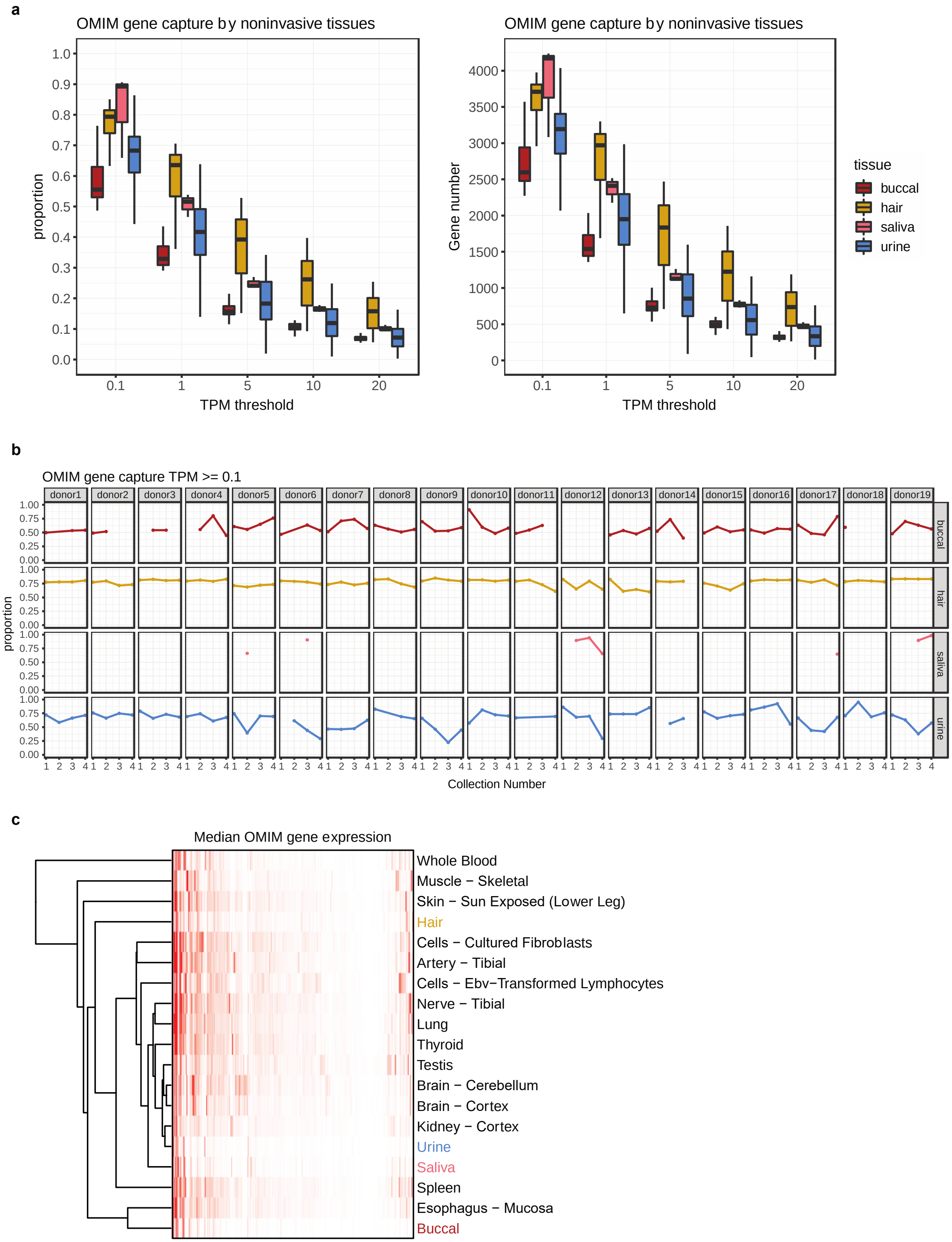
a
OMIM gene capture by noninvasive tissues
proportion
TPM threshold
0.1
1
5
10
20
OMIM gene capture by noninvasive tissues
Gene number
TPM threshold
tissue
buccal
hair
saliva
urine
b
OMIM gene capture TPM >= 0.1
donor1
donor2
donor3
donor4
donor5
donor6
donor7
donor8
donor9
donor10
donor11
donor12
donor13
donor14
donor15
donor16
donor17
donor18
donor19
buccal
hair
saliva
urine
proportion
Collection Number
c
Median OMIM gene expression
Whole Blood
Muscle − Skeletal
Skin − Sun Exposed (Lower Leg)
Hair
Cells − Cultured Fibroblasts
Artery − Tibial
Cells − Ebv−Transformed Lymphocytes
Nerve − Tibial
Lung
Thyroid
Testis
Brain − Cerebellum
Brain − Cortex
Kidney − Cortex
Urine
Saliva
Spleen
Esophagus − Mucosa
Buccal

**Supplementary Figure 13. a.** Proportion and total OMIM gene capture per noninvasive tissue type, depending on minimum median TPM threshold. **b.** Proportion of OMIM gene capture per collection, donor, and tissue using a minimum expression threshold of 0.1 TPMs. **c.** Clustering of noninvasive and select GTEx tissues based on median OMIM gene expression.

**Supplementary Table 1.** Breakdown of reaction cost (as of August 2022) per reagent using our in-house method (Loseq) versus TruSeq Stranded mRNA Library Prep and Takara SMART-seq V4 commercial kits. The cost of Ampure XP beads (cat# A63881) is excluded, but it is notably lower per reaction for Loseq and SMART-seq preparations due to smaller volume requirements.

| Catalog Number | Reagent Name | Cost per reaction |
|---|---|---|
| Loseq | | |
| 50-196-5299 | KAPA HiFi HS reaction mix | $1.38 |
| 30281-2 | NxGen® RNAse Inhibitor | $0.69 |
| EP0752 | Maxima H Minus RT | $0.48 |
| R0192 | dNTP set (10mM each) | $0.14 |
| | 3' RT Primer (anchor) HPLC purified | $0.28 |
| | drop-TSO HPLC Purification | $0.09 |
| | drop-PCR HPLC Purification | $0.04 |
| 20027213 | Nextera UD indexes | $1.75 |
| FC-131-1096 | Nextera XT DNA Library Preparation Kit | $8.95 |
| 61012 | Dynabeads mRNA Direct Purification kit | $3.75 |
| AM8170G | DNase I Buffer (10x) | $0.12 |
| AM2224 | DNase I (RNase-free) | $0.03 |
| Illumina | | |
| 20020595 | TruSeq Stranded mRNA | $96.46 |
| 20020591 | TruSeq UD indexes | $5.71 |
| SMART-seq | | |
| 634891 | TakaraBio SMART-seq V4 | $55.92 |

**Supplementary Table 2.** Comparison of total cost per reaction by library preparation. Loseq percent cost reduction calculated as: (1-(Loseq/Commercial kit))*100

| Library Preparation | Total Cost per Reaction | Loseq Cost Reduction |
|---|---|---|
| Loseq | $17.70 | |
| Illumina | $102.17 | 82.68% (-$84.47) |
| SMART-seq | $55.92 | 68.35% (-$38.22) |
